# Genome-wide screen of *Mycobacterium tuberculosis-*infected macrophages identified the GID/CTLH complex as a determinant of intracellular bacterial growth

**DOI:** 10.1101/2024.05.06.592714

**Authors:** Nelson V. Simwela, Luana Johnston, Paulina Pavinski Bitar, Eleni Jaecklein, Craig Altier, Christopher M. Sassetti, David G. Russell

**Affiliations:** Department of Microbiology and Immunology, College of Veterinary Medicine, Cornell University, Ithaca, New York, USA; Department of Population Medicine and Diagnostic Sciences, Cornell University, Ithaca, New York, USA; Department of Microbiology and Physiological Systems, UMass Chan Medical School, Worcester, Massachusetts, USA

**Keywords:** CRISPR genetic screen, *Mycobacterium tuberculosis*, macrophage cytotoxicity, GID/CTLH complex, autophagy, nutritional immunity

## Abstract

The eukaryotic GID/CTLH complex is a highly conserved E3 ubiquitin ligase involved in a broad range of biological processes. However, a role of this complex in host antimicrobial defenses has not been described. We exploited *Mycobacterium tuberculosis* (*Mtb*) induced cytotoxicity in macrophages in a FACS based CRISPR genetic screen to identify host determinants of intracellular *Mtb* growth restriction. Our screen identified 5 (*GID8*, *YPEL5*, *WDR26*, *UBE2H*, *MAEA*) of the 10 predicted members of the GID/CTLH complex as determinants of intracellular growth of both *Mtb* and *Salmonella* serovar Typhimurium. We show that the antimicrobial properties of the GID/CTLH complex knockdown macrophages are mediated by enhanced GABAergic signaling, activated AMPK, increased autophagic flux and resistance to cell death. Meanwhile, *Mtb* isolated from GID/CTLH knockdown macrophages are nutritionally starved and oxidatively stressed. Our study identifies the GID/CTLH complex activity as broadly suppressive of host antimicrobial responses against intracellular bacterial infections.

**Graphical abstract:** 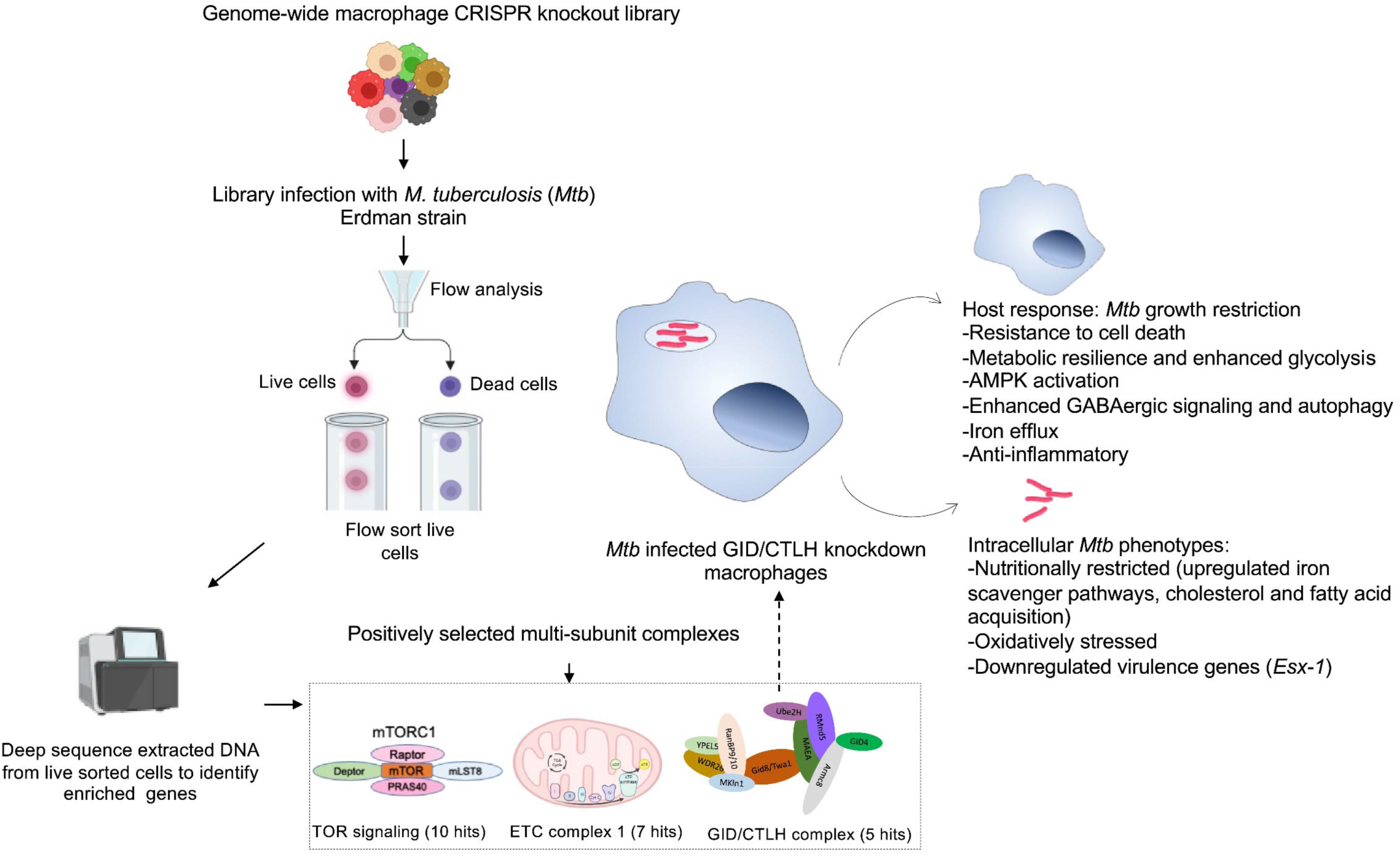

## Introduction

*Mycobacterium tuberculosis* (*Mtb*) is an obligate intracellular pathogen that causes tuberculosis (TB), the leading cause of global mortality due to a single infectious disease.^1^ At the onset of infection, *Mtb* is inhaled into the airways and phagocytosed by resident alveolar macrophages. This initial interaction triggers recruitment of monocyte-derived macrophages, which gradually become the more abundant host macrophage population as the infection becomes more established. It is estimated that only 5-10% of *Mtb* infected individuals will progress to active TB disease.^1^ Whether the non-progressors remain latently infected, or progress to clear the infection is a subject of on-going debate. *Mtb* has evolved to survive and grow within the macrophage through the subversion of the normal anti-microbial properties of the phagocyte. In brief, the bacterium blocks normal phagosomal acidification and phagosome/lysosome fusion.^2–5^ Inside the macrophage, the bacterium also relies on access to host fatty acids and cholesterol to fuel its biosynthetic demands.^6^ The outcome of this interplay is strongly influenced by both the macrophage lineages involved and the development of an acquired immune response. It is now known that lung macrophages comprise of two separate lineages; tissue resident alveolar macrophages that are fetal stem cell-derived and populate the lung during embryogenesis, and interstitial macrophages that are derived from blood monocytes and are recruited to the lung upon insult or infection.^7,8^ While resident alveolar macrophages are more permissive to *Mtb* growth, recruited interstitial macrophages are natively hostile to the intracellular resident bacteria.^9^ Eventually, *Mtb* parasitization of the macrophage usually ends in the death of the infected phagocyte. However, the route to cell death also plays a major impact on bacterial survival. If the infected macrophage undergoes apoptosis, *Mtb* will likely be killed, either in the apoptotic body or as a result of efferocytosis.^10,11^ In contrast, macrophage death via necrosis supports *Mtb* growth and aids in bacterial spread.^12,13^ One of the most significant determinants of *Mtb* fate inside macrophages is its ability to perforate the phagosome by an *EsX-1* type VII secretion system.^14,15^ Phagosomal damage releases *Mtb* effectors into the host cytosol that suppress apoptosis and drive necrosis.^12,16^ Through the action of *EsX-1*, *Mtb* also inhibits macrophage antimicrobial defenses such as autophagy and inflammasome mediated production of interleukin-1β (IL-1β).^17,18^

The route to macrophage cell death is clearly important to *Mtb*’s success as a pathogen. Two groups have conducted genome-wide CRISPR/Cas9 screens targeting *Mtb* induced macrophage cell death pathways. Zhang and colleagues screened RAW264.7 cells and identified multiple genes in the type 1 interferon signaling pathway that, when disrupted, conferred protection against *Mtb*-induced cell death.^19^ Similarly, Lai et al. performed a whole genome screen on THP1 cells infected with *M. bovis* BCG and also identified hits in the type 1 interferon pathway as having protective effects on infected cells.^20^ Both of these screens used the diminished death of infected cells to select for genes that, when deleted, provided the cells with some degree of protection. In this current study, we used a similar CRISPR screening approach but emphasized a different selection mechanism. We hypothesized that improved cell survival could also be a result of mutations that restrict bacterial growth, which has the potential to identify novel host pathways of bacterial control. We performed a genome wide CRISPR/Cas9 screen on primary Hoxb8 conditionally immortalized murine bone marrow macrophages and identified the mammalian Glucose Induced Degradation (GID) / C-terminal to LisH (CTLH) complex as a strong determinant of intracellular growth of *Mtb* in macrophages. We show that knockdown of individual components of the GID/CTLH complex is strongly restrictive to the intracellular growth of *Mtb* and *S.* Typhimurium. Mechanistically, knockdown of the GID/CTLH complex promotes antimicrobial responses in macrophages through an enhanced autophagic response, increased metabolic resilience and resistance to cell death. GID/CTLH knockdown macrophages are also anti-inflammatory and at the same time restrict intracellular *Mtb* access to essential nutrients. Our work demonstrates that the GID/CTLH complex activity broadly suppresses the macrophage’s antimicrobial responses and therefore represents a tractable target for host directed therapies (HDTs) in TB control strategies.

## Results

### FACS based macrophage survival CRISPR screen to identify intracellular determinants of Mtb replication

Infection of macrophages with virulent *Mtb* strains induces necrotic cell death,^12,13,21^ a virulence mechanism that enables the bacteria to escape the phagosome and spread. To identify genes that contribute to improved macrophage survival after *Mtb* infection, we developed a FACS based CRISPR screen in murine macrophages derived from estradiol responsive Hoxb8 Cas9^+^ conditionally immortalized myeloid precursors^22^ using cytotoxicity as a selection strategy, anticipating that restriction of *Mtb* growth would support better cell survival. We first optimized *Mtb* cytotoxicity in Hoxb8 bone marrow derived macrophages (hBMDMs) at different multiplicity of infections (MOIs). Infection of hBMDMs with the virulent *Mtb Erdman* strain induced ∼50% macrophage cytotoxicity at MOI 1 and >75% cytotoxicity at MOI 3 after 4 days of infection as measured by the lactate dehydrogenase (LDH) release assay (Fig. 1A). Similar levels of cytotoxicity were evident when we analyzed the infected macrophages by flow cytometry after staining with the live dead viability dye (Fig.S1B). As further evidence of bacteria release from dying macrophages, an MOI dependent increase in *Mtb* colony forming units (CFUs) was observed in supernatants from the infected macrophage cultures (Fig. 1B). We therefore chose a 4-day infection with the *Mtb Erdma*n strain at MOI 1 as a selection pressure for our CRISPR screen as it allowed for sufficient proportions of surviving macrophages (Fig. S1B) which can be flow sorted for downstream library preparation and sequencing. We then transduced Hoxb8 Cas9^+^ myeloid progenitors with the Brie knockout lentiviral single guide RNA (sgRNA) library^23^ that targets ∼19,674 of mouse protein coding genes with 4 sgRNAs/gene. hBMDMs knockout libraries were differentiated from the Hoxb8 Cas9^+^ myeloid progenitor library (Fig. S1A) and infected with the *Mtb* Erdman strain at MOI 1 for 4 days. Surviving macrophages based on live dead staining were flow sorted and sequenced to identify candidate genes (Fig. 1C). We carried out the screen in 3 independent replicates and identified hits by comparing sgRNA read counts in the sample groups to the input unperturbed hBMDM library using the MAGeCK pipeline.^24–26^

**Fig.1.**
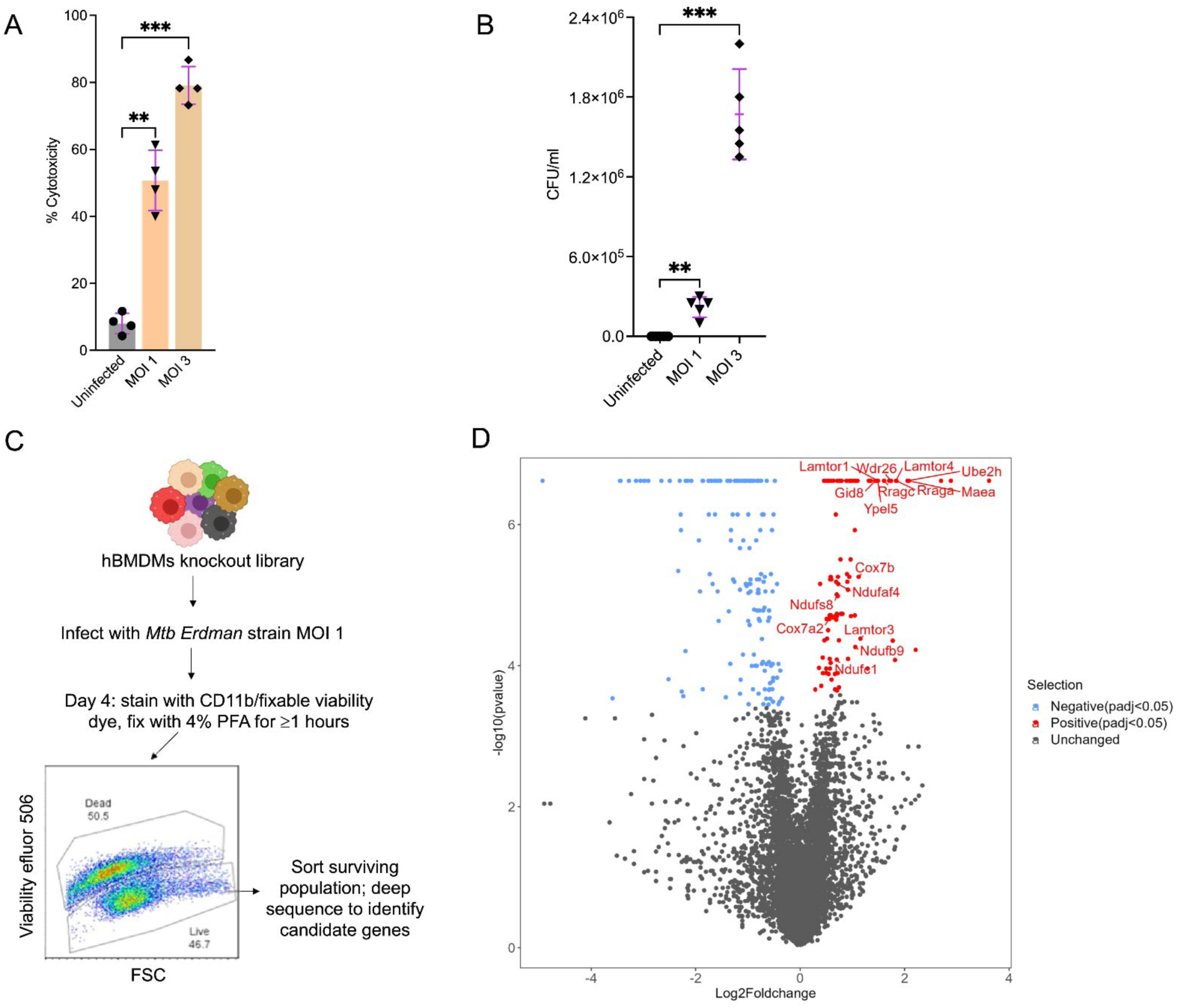
Genome wide CRISPR screen to identify host determinants of tuberculosis restriction in macrophages. (A) Mycobacterium tuberculosis (Mtb) induced cytotoxicity in Hoxb8 bone marrow derived macrophages (hBMDMs) at the indicated multiplicity of infections (MOI) 4 days post infection. Cytotoxicity was quantified by measuring the activity of released lactate dehydrogenase (LDH) in cell supernatants as compared to 100% LDH activity in detergent lysed cells; n=3. **P < 0.01; ***P < 0.001. (B) A further quantification of Mtb induced cytotoxicity in macrophages at indicated MOIs by plating colony forming units (CFUs) in cell supernatants from A; n=3. **P < 0.01; ***P < 0.001. (C) Schematic workflow for performing CRISPR screen to identify host determinants of Mtb restriction in macrophages. (D) Volcano plot showing hits from the screen. For each gene (represented by dots), enrichment or depletion is shown in the x-axis as log2 fold change while the y-axis shows the corresponding-log10 p value. Significantly enriched and depleted hits are shown in red and blue colors respectively while unchanged hits are in grey. For significant hits, an adjusted p value of <0.05 was used as a cutoff. Screens were carried out in three independent replicates.

We identified 259 genes whose knockdown modulated hBMDMs responses to *Mtb* infection based on a false discovery rate of <0.05 (Fig. 1D, Table.S1). 104 genes were significantly enriched (conferred relative protection to *Mtb* induced cytotoxicity) while 155 genes were significantly depleted. The highest scoring protective hits belonged to the TOR signaling pathway (Fig. 1D, Table S1, S2) which has already been implicated in macrophage control of *Mtb*.^27^ All the enriched 10 genes in the TOR pathway (*RRAGC*, *LAMTOR1*, *RRAGA*, *TSC1*, *FLCN*, *TSC2*, *LAMTOR4*, *LAMTOR3*, *TBC1D7*, *RPS6KA1*) are part of the mTOR signaling cascade which plays a crucial role in mammalian cell growth and nutrient signaling.^28^ These hits are consistent with the literature that mTOR inhibition is known to activate autophagy, a host response that restricts intracellular *Mtb* growth in macrophages.^29^ Our hits also included a number of genes and pathways that could impact negatively on *Mtb* growth (Fig. 1D, Table S2) such as the mitochondrial complex 1 (*NUBPL*, *TMEM126B*, *NDUFAF4*, *NDUFS8*, *ACAD9*, *NDUFB9*, *NDUFC1*), iron-sulfur cluster binding, protein dephosphorylation, regulation of GTPases activity and lysosomal functions. Mitochondrial complex 1 is a target of metformin, a promising HDT against *Mtb* ^30^ while disruption of iron homeostasis in macrophages by knockdown of the iron-sulfur clusters could be restrictive to *Mtb* growth by limiting the bacteria’s access to iron.^31^ We also identified 5 of the 10 predicted members of the highly conserved mammalian multi-subunit GID/CTLH complex^32,33^ (*MAEA, WDR26*, *UBE2H*, *YPEL5*, *GID8*) (Fig. 1D, Table S1) as potential determinants of macrophage resistance to *Mtb* driven cell death. Interestingly, 2 of these hits (*WDR26*, *YPEL5*) were previously identified in a related CRISPR screen in THP-1 macrophages using *M. bovis* BCG.^20^

### Knockdown of the GID/CTLH complex in macrophages results in strong intracellular bacteria growth restriction and confers high level resistance to *Mtb* induced cell cytotoxicity

The GID/CTLH complex was first identified in yeast as a multi-subunit E3 ligase that targets surplus gluconeogenic enzymes for proteasomal degradation when glucose starved cells are re-supplied with the substrate.^34,35^ In animals, the complex appears to fulfill other biological functions like cell proliferation and survival, cell migration and adhesion, erythropoiesis and neurodegenerative diseases.^36^ In immunity, some members of the complex (*RANBP9*) have been implicated in antigen processing, efferocytosis and anti-inflammatory activities.^37,38^ We generated *de novo* knockdowns in Hoxb8 Cas9^+^ myeloid progenitors for the 5 GID/CTLH hits identified in our screen (*MAEA, WDR26*, *UBE2H*, *YPEL5*, *GID8*) which represent ∼50% of the predicted functional members of the complex in humans and mice.^32^ We targeted each gene with 2 sgRNAs and were able to achieve >85% knockdown efficiencies (Table S3) as analyzed by the Inference of CRISPR Edits (ICE) tool.^39^ Similar levels of protein knockdown were also confirmed by western blot analysis for all the 5 genes in hBMDMs differentiated from the Hoxb8 Cas9^+^ mutants (Fig. S2). We then infected the GID/CTLH knockdown hBMDMs differentiated from the Hoxb8 Cas9^+^ mutants with the *Mtb Erdman* strain at MOI 0.4 for 4 days to determine the intracellular growth replication rates of *Mtb* in these macrophages. A strong growth restriction was observed in all the 5 GID/CTLH mutants as compared to macrophages transduced with a non-targeting scramble sgRNA control (Fig. 2A). The growth restriction phenotypes were not due to differences in bacteria uptake or phagocytosis defects as there was no significant differences when we plated CFUs 3 hours post infection (Fig. 2A). Similar intracellular growth restriction phenotypes were observed when we infected a selected mutants (*MAEA*, *GID8*, *WDR26*) with the *Mtb Erdman*-Lux strain and monitored bacterial growth by luciferase expression (Fig. 2B).

**Fig.2.**
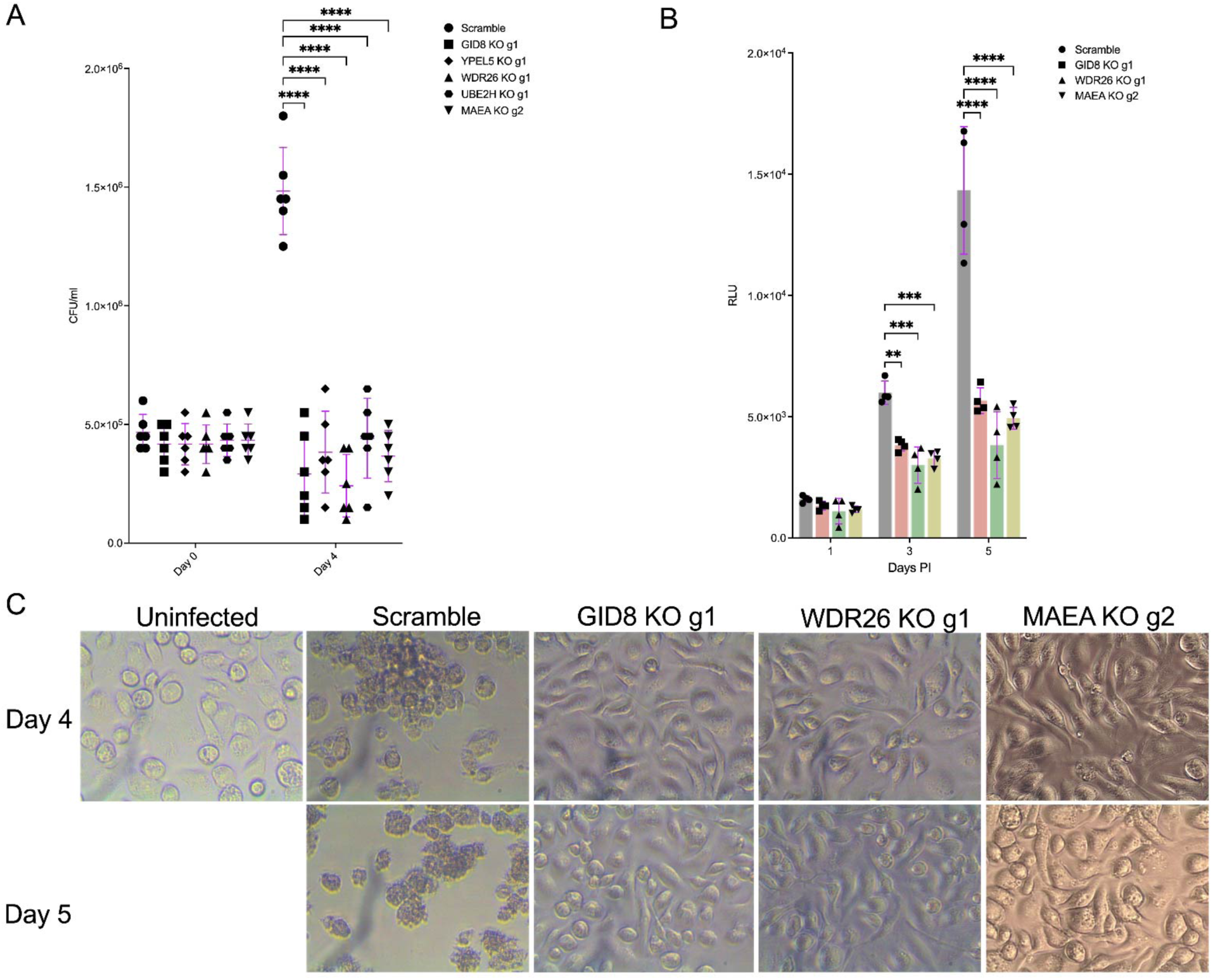
Macrophages with knockdown of the GID/CTLH complex are strongly restrictive to *Mtb* intracellular growth and are resistant to *Mtb* induced cell death. (A) Wild type (scramble) or indicated GID/CTLH knockdown hBMDMs were infected with *Mtb Erdman* strain at MOI 0.4. CFUs were plated on day 0 (3 hours post infection) and on Day 4 to determine intracellular *Mtb* replication rates; n=6. ****P < 0.001. (B) Quantification of *Mtb* replication in scramble or *GID8, MAEA* and *WDR26* knockdown hBMDMs using the *Mtb Erdman*-Lux strain. hBMDMs were infected at MOI 0.5 following which luciferase measurements were taken on the indicated days; n=4. **P < 0.01; ***P < 0.001; ****P < 0.0001. (C) Microscopy micrographs of scramble or *GID8*, *MAEA* and *WDR26* knockdown hBMDMs infected with *Mtb Erdman* strain at MOI 1 taken on day 4 and 5 post infection.

We also analyzed the growth restriction phenotypes of the GID/CTLH knockdown hBMDMs using another intracellular bacterium, *Salmonella enterica* serovar Typhimurium (*S.* Typhimurium), in the gentamicin protection assay.^40^ *MAEA* and *GID8* knockdown hBMDMs were infected with the wild type *S.* Typhimurium ATCC 14028s strain at MOI 10 and CFUs were plated 4 and 18 hours post infection. Again, we observed a strong intracellular growth restriction of *S.* Typhimurium in these mutant macrophages, compared to scramble sgRNAs (Fig. S3A). The growth restriction phenotypes were also not due to bacterial uptake differences as similar bacteria numbers were evident in scramble and mutant macrophages infected with the isogenic BFP-expressing *S.* Typhimurium strain, *phoN*::BFP,^41^ as analyzed by confocal microscopy (Fig. S3B). To further confirm that GID/CTLH knockdown growth restriction phenotypes in macrophages were also applicable in human cells, we generated *GID8* and *MAEA* knockdowns in primary human monocyte derived macrophages (hMDMs) by directly electroporating CRISPR ribonucleoproteins (CRISPR RNPs) in monocytes.^42^ We targeted each gene with at least 2 sgRNAs and were able to achieve 50-70% knockdown efficiency with some sgRNAs (Fig. S4A, Table S3). More importantly, we observed *Mtb* growth restrictions levels that were dependent on the degree of protein knockdown in these hMDM mutants by both CFUs and luciferase readouts (Fig. S4B, S4C). These data illustrated the functional conservation of the GID/CTLH complex in mediating intracellular growth restriction of bacteria in both mice and humans. We also analyzed GID/CTLH knockdown hBMDMs ability to resist cell death upon infection with *Mtb* by infecting the mutant macrophages with the *Mtb Erdman* strain at MOI 1 and monitoring macrophage survival by recording microscopy micrographs on day 4 and 5; infection conditions that mimicked our CRISPR screen. *Mtb* infection induced macrophage death that resulted in about half of the cells dying by day 4 in the scramble macrophages and almost all the cells appeared necrotic by day 5 (Fig. 2C). However, GID/CTLH knockdown macrophages were highly resistant to cell death with 100% survival rates on day 4 and 5. Together, these data suggested that knockdown of the GID/CTLH complex does not just restrict the intracellular growth of *Mtb* in macrophages but also inhibits necrotic cell death programs triggered by the bacteria.

### *Mtb* infected GID/CTLH knockdown macrophages are metabolically resilient and more glycolytic

In yeast, the GID/CTLH complex is directly involved in glucose metabolism by regulating gluconeogenesis.^34^ Even though some studies have shown that this complex is dispensable in the regulation of carbohydrate metabolism in human cells,^32^ others have identified glycolytic enzymes L-lactate dehydrogenase A chain (LDHA) and pyruvate kinase M1/2 (PKM) as direct targets of GID/CTLH ubiquitination.^43^ In fact, depletion of the GID/CTLH complex member *RMND5A* reduces polyubiquitination of LDHA and PKM and makes the cells more glycolytic.^43^ Moreover, the GID/CTLH complex (member *RMND5A*) appear to regulate cell energy homeostasis by negatively modulating the activity of the AMP-activated protein kinase (AMPK) in *C. elegans.* ^44^ We therefore profiled the metabolic state of GID/CTLH knockdown hBMDMs in uninfected or *Mtb* infected conditions. We first checked the relative mRNA expression of the glycolytic enzyme hexokinase 2 (*Hk2*) and the gluconeogenic enzyme fructose-1,6-bisphosphatase 1 (*FBP1*) in these GID/CTLH knockdown hBMDMs. We observed a significant increase in mRNA expression of *HK2* in uninfected GID/CTLH knockdown hBMDMs as compared to scramble (Fig. S5A). *Mtb* infection increased the expression of *HK2* in GID/CTLH knockdown hBMDMs 4 hours post infection as compared to scramble, but these responses resolved after 24 hours of infection (Fig. S5A). This is in line with a biphasic metabolic profile of macrophage responses to *Mtb* infection which is characterized by an early upregulation of glycolytic genes in the first 4-8 hours of infection that diminishes 24-48 hours later.^45^ We, however, did not observe any differences in *FBP1* expression in uninfected or *Mtb* infected GID/CTLH knockdown hBMDMs (Fig. S5A). These results suggest that GID/CTLH knockdown in hBMDMs does not impact gluconeogenesis (at least at transcript level) but programs the cells to a more glycolytic state, which is consistent with previous observations.^43^ To further confirm these phenotypes, we analyzed the metabolic status of GID/CTLH knockdown hBMDMs by Seahorse extracellular flux analyses using the Agilent Mito and Glycolysis Stress Test kits. Compared to scramble sgRNA, uninfected GID/CTLH knockdown hBMDMs displayed reduced basal oxygen consumption rates (OCRs) and spare respiratory capacity (SRC) (Fig. S5B) which suggested a reduction in mitochondrial activities and oxidative phosphorylation as a consequence of GID/CTLH knockdown. In agreement with our qPCR data (Fig. S5A), we also observed increased glycolytic flux in uninfected GID/CTLH knockdown hBMDMs as evidenced by high basal extracellular acidification rates (ECAR) and spare glycolytic capacity (SGC) (Fig. S5C). Under basal conditions, *Mtb* infection of wild type macrophages increases glycolytic rates that are required to control bacterial replication^46^ while at the same time decelerates mitochondrial respiration.^47^ Indeed, when we infected scramble macrophages with *Mtb* for 24 hours, OCR rates collapsed to almost baseline levels (Fig. 3A, 3B) as compared to uninfected (Fig. S5A). However, *Mtb* infected GID/CTLH knockdown macrophages were more resilient as they maintained significantly higher basal OCRs and SRCs as compared to scramble sgRNA (Fig. 3A, 3B). Glycolytic rates in *Mtb* infected GID/CTLH knockdown hBMDMs were also significantly higher as compared to scramble sgRNA (Fig. 3C,3D) albeit at an even higher rate than the uninfected (Fig. S5C). Together, qPCR and extracellular flux analysis data showed that knockdown of the GID/CTLH complex in macrophages impairs mitochondrial respiration and proportionally increases flux through glycolysis. However, GID/CTLH knockdown hBMDMs are able to withstand an *Mtb* induced bioenergetic shutdown^47^ despite displaying reduced mitochondrial activities in uninfected conditions. Our metabolic analyses also pointed to a state of energy stress in GID/CTLH knockdown hBMDMs which could be regulated, in part, by AMPK, a master regulator of cell energy homeostasis.^48^ Given that AMPK can be modulated (negatively) by the GID/CTLH complex, at least in nematodes,^44^ we also checked AMPK activity in uninfected and *Mtb* infected GID/CTLH knockdown macrophages by monitoring AMPK phosphorylation status in western blot analyses. Consistent with our metabolic profiling data, we observed significant AMPK phosphorylation in both uninfected and *Mtb* infected GID/CTLH knockdown macrophages while the levels of total AMPK were mostly unchanged (Fig. 3E).

**Fig.3.**
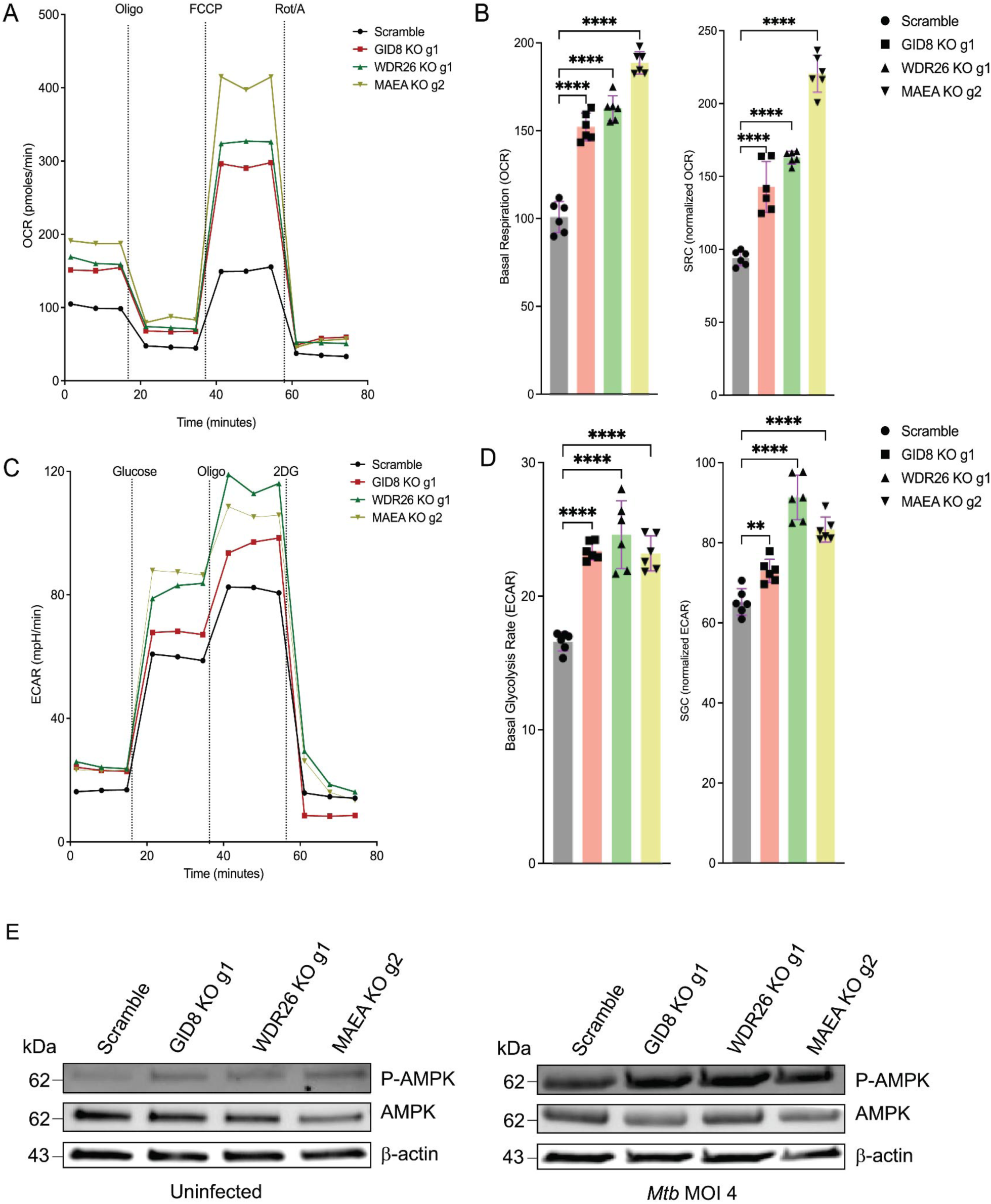
*Mtb* infected GID/CTLH knockdown macrophages have altered metabolism and are strongly glycolytic. (A) Flux analyses of scramble or *GID8*, *MAEA* and *WDR26* knockdown hBMDMs infected with *Mtb Erdman* strain at MOI 1 24 hours post infection. Oxygen consumption rates (OCR) were measured using the Cell Mito Stress Test Kit (Agilent). Oligo, oligomycin; FCCP, fluoro-carbonyl cyanide phenylhydrazone; Rot/A, rotenone and antimycin A. (B) Comparison of basal respiration and spare respiratory capacity (SRC) from A. SRC was calculated by subtracting the normalized maximal OCR from basal OCR; n=3 (2 technical replicates per repeat). ****P < 0.0001. (C) Extracellular acidification rates (ECARs) of scramble or GID8, *MAEA* and *WDR26* knockdown hBMDMs infected with *Mtb* as in A. ECARs were measured using the Agilent Seahorse Glycolysis Stress Test kit. 2DG, 2-Deoxy-D-glucose. (D) Comparison of basal glycolysis and spare glycolytic capacity (SGC) in the indicated hBMDM knockdowns. SGC was calculated as SRC above. n=3 (2 technical replicates per repeat). **P < 0.01; ****P < 0.0001. (E) Western blot analysis of AMPK phosphorylation in scramble or *GID8*, *MAEA* and *WDR26* knockdown hBMDMs uninfected or infected with *Mtb* at MOI 4.

### Dual RNA sequencing identify GABAergic signaling and autophagy as major host determinants of *Mtb* restriction in GID/CTLH knockdown hBMDMs

To further identify effectors that mediate *Mtb* growth restriction in GID/CTLH knockdown hBMDMs, we performed dual RNA sequencing (dual RNA-seq) of *Mtb* infected knockdown macrophages to capture both host and bacterial transcriptomes.^49^ Scramble sgRNAs or GID/CTLH knockdown hBMDMs (*GID8, MAEA, WDR26*) were infected with the *Mtb smyc’::mCherry* strain for 4 days. On day 4, we flow sorted *Mtb* infected macrophages based on mCherry positivity and extracted RNA for dual RNA-seq.^49^ Principal component analysis (PCA) of the macrophage transcriptomes revealed unique clustering of *GID8, MAEA* and *WDR26* knockdown hBMDMs which despite being slightly distant from each other, separated close together in the first principal component as compared to scramble sgRNA (Fig. 4A). Using an adjusted p value of < 0.05 and an absolute log2 fold change of > 0.5, we identified 1911 upregulated and 918 downregulated genes in *GID8* mutant macrophages, 2106 upregulated and 940 downregulated genes in *MAEA* mutant macrophages and 2890 upregulated and 1845 downregulated genes in *WDR26* mutant macrophages (Table S4). Due to the similar separation of *GID8, MAEA* and *WDR26* in our PCA analysis (Fig. 4A), we plotted a Venn diagram to identify genes which were commonly differentially expressed (DE) between the 3 mutant macrophage populations. Indeed, a majority of genes were commonly DE (1267 upregulated, 421 downregulated) between *GID8, MAEA* and *WDR26 Mtb* infected hBMDMs (Fig. 4B, Table S4). Among the highly upregulated genes in the commonly DE gene set were those involved in macrophage effector functions that are known to control *Mtb* replication such as the NADPH oxidase (*NOX3*) and autophagy (*ATG9B, TRIM2, TRIM7 and TRIM16*) (Table S4). Ferroportin 1 (*SLC40A1*), the only known membrane protein to transport iron out of the cells,^50^ was also among the most significantly upregulated genes in *GID8, MAEA* and *WDR26 Mtb* infected hBMDMs (Table S4). This suggested that GID/CTLH knockdown in hBMDMs could restrict *Mtb* growth by nutrient (iron) limitation, enhancing autophagy and increasing production of reactive oxygen species (ROS).

**Fig.4.**
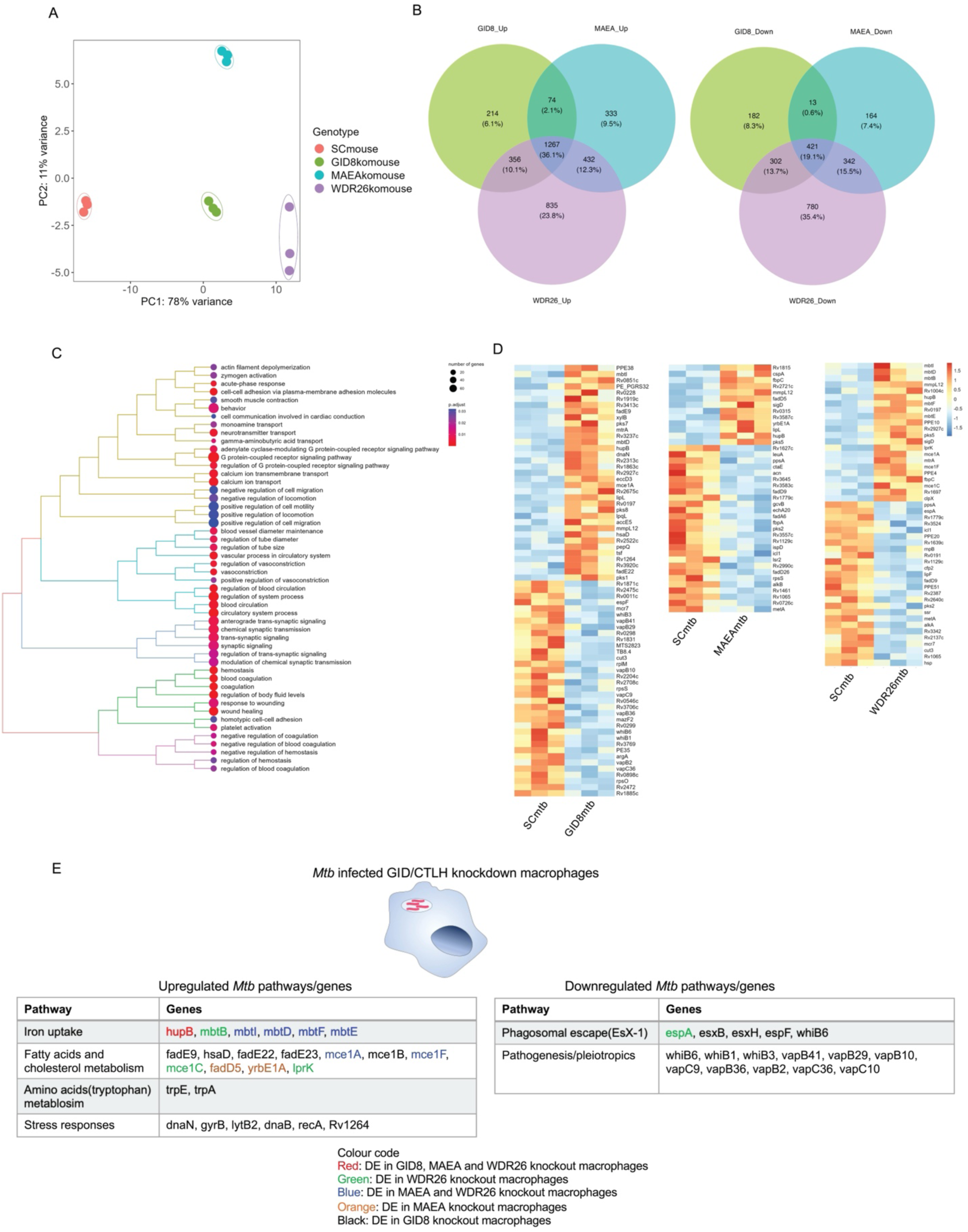
Host and bacterial transcriptional responses to identify determinants of *Mtb* restriction in GID/CTLH knockout macrophages. (A) Principal component analysis (PCA) of scramble or *GID8*, *MAEA* and *WDR26* knockdown hBMDMs transcriptomes infected with the *Mtb smyc’::mCherry* strain at MOI 0.5 4 days post infection (B) Venn diagrams of upregulated and downregulated gene sets (Table. S4) in *GID8*, *MAEA* and *WDR26* knockdown hBMDMs as compared to scramble showing overlapping genes. Differentially expressed genes cutoff; abs (log2 fold change) > 0.5, adjusted p-value < 0.05. (C) Tree plot of top 50 enriched gene ontology terms (biological process) amongst the 1267 commonly upregulated genes in *GID8*, *MAEA* and *WDR26* knockdown hBMDMs (D) Heatmaps of the top differentially expressed genes in *Mtb* isolated from *GID8* (70 genes), *MAEA* (40 genes) and *WDR26* (50 genes) knockdown hBMDMs (E) Schematic of some of the differentially expressed genes in *Mtb* isolated from *GID8*, *MAEA* and *WDR26* knockdown hBMDMs categorized into pathways

To gain more insights into biological pathways that could mediate *Mtb* growth restriction in GID/CTLH knockdown macrophages, we performed pathway enrichment analysis^51^ of the *GID8, MAEA* and *WDR26* commonly DE genes (Table S4). In the upregulated gene set, we identified 60 biological processes (BP), 30 cellular components (CC) and 27 molecular functions (MF) which were enriched based on an adjusted p value of < 0.05 (Fig. 4C, Table S5). Amongst the commonly downregulated DE genes, 332 BP, 16 CC and 44 MF were enriched (Fig. S6, Table S5). Of the top enriched BPs in the commonly upregulated genes (Fig. 4C), the most enriched pathways included those involved in Ca^2+^ and G protein receptor signaling, effector processes which are known to play important roles in macrophage responses against *Mtb*.^52,53^ Of note, γ-Aminobutyric acid (GABA) and neurotransmitter transport were, similarly, amongst the most enriched pathways in the GID/CTLH knockdown upregulated genes (Fig. 4C, Table S5). Within the GID/CTLH complex, only 1 member (Muskelin, *MKLN1*) has been shown to be involved in GABA receptor trafficking and internalization.^54^ We, however, identified 6 genes in the GABAergic signaling pathway (*GABBR1*, *GABRB2*, *GABARAPL1*, *SLC32A1*, *SLC6A13*, *SLC6A12*) which were upregulated in all the 3 *Mtb* infected GID/CTLH knockdown mutants; *GID8*, *MAEA* and *WDR26* (Table S4, S5). In fact, *GABBR1* and *GABRB2* were amongst the most significantly upregulated genes in the GID/CTLH knockdown macrophages with up to 350-fold upregulation as compared to scramble sgRNA (Table S4). Only recently, GABAergic signaling has been linked to intracellular macrophage control of *Mtb* and *S.* Typhimurium. Treatment of macrophages with GABA or GABAergic drugs enhances the antimicrobial properties of macrophages against intracellular bacterial infections by promoting phagosome maturation and autophagy.^55^ Moreover, the apparent GABAergic protective responses in BMDM macrophages were driven, in significant part, by increase in Ca^2+^ signaling.^55^ Taken together, our pathway enrichment data suggested that GID/CTLH knockdown in hBMDMs increases GABA and Ca^2+^ signaling which may, similarly, promote macrophage control of *Mtb* by promoting autophagy.^55^ A further consequence of GABA treatment in *Mtb* infected bMDMs and in infected mice lungs *in vivo* was the inhibition of proinflammatory markers such as Tnf and Il-6.^55^ Similarly, proinflammatory pathways were the most significantly enriched pathways in our GID/CTLH commonly downregulated genes (Fig. S6A, Table S5). These included pathways involved in IL-1β production, antigen presentation, toll receptor signaling and cellular responses to Tnf. Our further analysis of the RNA seq data did indeed show a significant downregulation of IL-1β and Tnf transcripts in all the GID/CTLH knockdown macrophages infected with *Mtb* on day 4 (Fig. S6B). Consistent with the known role of the GID/CTLH complex in cell proliferation,^36^ pathways related to DNA replication were also among the most enriched in the downregulated gene set (Fig. S6A). We also observed a significant downregulation of pathways related to cell death programs and apoptosis signaling (Table S5) in GID/CTLH knockdown hBMDMs which would possibly explain the increased resistance of these macrophages to *Mtb* induced cytotoxicity (Fig. 2C).

### Nutritional stress is the main transcriptome signature of intracellular *Mtb* in GID/CTLH knockdown hBMDMs

To assess the impact of host stressors on *Mtb*, we analyzed parallel transcriptomes of intracellular *Mtb* (Fig. 4A). Using an adjusted p value of < 0.05 and an absolute log2 fold change of > 0.3, we identified 168 *Mtb* genes (87 upregulated, 81 downregulated) in *GID8* knockdown hBMDMs, 40 *Mtb* genes (13 upregulated, 27 downregulated) in *MAEA* knockdown hBMDMs and 46 *Mtb* genes (22 genes upregulated, 26 downregulated) in *WDR26* knockdown hBMDMs which were DE (Fig. 4D, Table S6). Interestingly, clusters of upregulated *Mtb* genes in these GID/CTLH knockdown hBMDMs were readily categorized into essential *Mtb* biological processes such as iron scavenging, cholesterol breakdown, fatty acid oxidation, amino acid metabolism and stress responses (Fig. 4E). *Mtb* utilizes Fe^3+^ iron specific siderophoroes, mycobactins and carboxymycobactins, to scavenge and acquire iron from the host for its nutritional requirements. *Mtb* mycobactins are organized into a cluster of *Mbt* (*MbtA*-*MbtJ*) and *Mbt-2* (*MbtK*-*MbtN)* genes.^56^ We identified 5 genes in the *Mbt* cluster (*MbtB*, *MbtI*, *MbtD*, *MbtE, MbtF*) which were upregulated in *Mtb* transcriptomes isolated from GID/CTLH knockdown hBMDMs (Fig. 4E, Table S6). A positive regulator of *Mbt* operonic gene expression, *HupB*,^57^ was also upregulated in all the 3 *Mtb* transcriptomes isolated from GID/CTLH knockdown macrophages (Fig. 4E, Table S6). These data suggested that GID/CTLH knockdown macrophages, in part, limit intracellular *Mtb* growth by reducing the bacteria’s access to iron. This is consistent with our host transcriptome data which showed a significant upregulation of the only known iron efflux transporter (*SLC40A1*) in GID/CTLH knockdown macrophages (Table S4). The data is also consistent with our previous findings which have shown that iron limitation is one of the prominent stresses experienced by *Mtb in vivo* in lung interstitial macrophages which are less permissive to *Mtb* growth^49^ and that similar *Mtb* restrictive phenotypes can be reproduced in human macrophages using iron chelating chemical inhibitors.^31^ In the meanwhile, *Mtb* nutrient limitation in GID/CTLH knockdown macrophages was not limited to iron as we also observed an upregulation of genes involved in beta oxidation of fatty acids (*FadE9*, *FadD5*, *FadE22*, *FadE23*),^58^ cholesterol breakdown (*HsaD*)^59^ and several members of the *Mce1* operon (*Mce1A*, *Mce1B*, *Mce1F*, *Mce1C*, *yrbE1A*, *IprK*) which is required for *Mtb* fatty acid import.^60,61^ Our previous work has demonstrated that limiting *Mtb*’s access to iron can upregulate cholesterol metabolism genes to re-balance impaired metabolic fluxes that arise due to diminished activity of iron dependent metabolic pathways.^31^ Even though upregulation of *Mtb* lipid metabolism genes in GID/CTLH knockdown macrophages could, indeed, be a compensatory mechanism to iron limitation,^31^ *Mtb* nutrient restriction in these macrophages appear to be more widespread as the bacteria also upregulates genes involved in tryptophan biosynthesis pathways (*TrpE*, *TrpA*) (Fig. 4E, Table S6). Tryptophan can serve as an alternative carbon source in these starved macrophage environments given that this amino acid is required for intracellular *Mtb* survival.^62^ GID/CTLH knockdown macrophages also appear to induce other non-nutritional stresses to intracellular *Mtb* as evidenced by the upregulation of genes required to possibly survive low PH in the phagosome (*Rv1264*).^63^ *Mtb* DNA damage and repair genes (*DnaN*, *DnaB*, *gyrB*, *recA*) (Fig. 4E) were also upregulated which is indicative of increased oxidative stress as we have previously observed in *Mtb* isolated from growth restrictive interstitial macrophages *in vivo*.^49^

More interestingly, *Mtb* in GID/CTLH knockdown macrophages downregulated major virulence genes including those belonging to the *EsX-1* type VII secretion system (Fig. 4E). We identified 4 genes in the *EsX-1* operon (*espA*, *esxB*, *esxH*, *espF*) which were significantly downregulated in *Mtb* isolated from GID/CTLH knockdown macrophages. A further 3 genes (*WhiB1, WhiB3, WhiB6*) belonging to the *WhiB* like family of transcription factors that play key roles in *mycobacteria* virulence and antibiotic resistance^64^ were also significantly downregulated (Fig. 4E, Table S6). Of note, *WhiB6* is one of the main regulators of *Esx-1* gene expression in *Mtb*.^65,66^ Central to the activity of all *WhiB* transcription factors are highly conserved cysteine bound iron-sulfur clusters that can act as reductive sinks to host nitric oxide or ROS produced during macrophage activation.^64^ In line with this, it has, indeed, been demonstrated that disruption of iron-sulfur clusters in *WhiB* transcription factors either by iron limitation or excessive nitrosative stress can trigger a significant reprogramming of gene expression in *Mtb*.^67^ Downregulation of *Mtb WhiB* genes in GID/CTLH knockdown macrophages could also, therefore, be a result of iron-limiting conditions in these mutant phagocytes which in case of *WhiB6* could trigger a feed forward effect to downregulate downstream *Esx-1* genes. As a consequence of *Esx-1* downregulation, *Mtb* resident in GID/CTLH knockdown macrophages could be attenuated and possibly trapped in phagosomes given the role of this secretion system in *Mtb* virulence and phagosomal escape.^17,68^ Several toxin-antitoxin (TA) genes were also downregulated in *Mtb* isolated from GID/CTLH knockdown macrophages (Fig. 4E, Table S6). Despite a reductive evolution in the *Mtb* genome, TA systems have been maintained to support *Mtb* replication, virulence and stress adaptation.^69^ Reduced expression of *Mtb* TA genes in GID/CTLH knockdown macrophages could thus be a further indication of shutdown in most bacterial virulence processes alongside the observed growth restriction phenotypes (Fig. 2A, 2B).

### GID/CTLH knockdown macrophages display increased GABAergic signaling, are anti-inflammatory and more autophagic

We followed up on our dual RNA-seq data (Fig. 4) to experimentally validate some of the host effectors that mediate *Mtb* growth restriction in GID/CTLH knockdown hBMDMs. We first checked the levels of GABA signaling in mutant macrophages by staining the cells for total endogenous GABA with an anti-GABA antibody. Murine BMDMs and lung macrophages express functional GABA in resting states^55^ and we indeed, similarly, observed expression of GABA in uninfected scramble hBMDMs when we analyzed anti-GABA-stained cells by confocal microscopy (Fig. S7A). Meanwhile, all the 3 uninfected GID/CTLH knockdown hBMDMs had significantly higher levels of GABA as compared to scramble (Fig. S7A, Fig. 5B). It was previously observed that *Mtb* infection of mouse BMDMs *in vitro* and *in vivo* significantly decreases GABA levels.^55^ In agreement with these previous findings, we also observed an almost shutdown of GABA expression in *Mtb* infected scramble hBMDMs (Fig. 5A). On the contrary, *Mtb* infection significantly amplified GABA levels in GID/CTLH knockdown hBMDMs to even greater levels than the uninfected (Fig. 5A, 5B). Increased GABAergic signaling can positively modulate intracellular Ca^2+^ mobilization.^70^ Moreover, Ca^2+^ signaling and transport were some of the enriched pathways in our GID/CTLH knockdown hBMDMs commonly upregulated genes (Fig. 4C). We therefore checked intracellular Ca^2+^ mobilization in GID/CTLH knockdown hBMDMs by loading the cells with a cell permeable fluorogenic calcium-binding dye (Fluoforte®) followed by quantification of emitted fluorescence post stimulation with a Ca^2+^ mobilization agonist, adenosine triphosphate (ATP). Compared to scramble sgRNA hBMDMs, uninfected GID/CTLH knockdown hBMDMs displayed increased intracellular calcium mobilization as evidenced by significantly higher normalized fluorescence 5 minutes after stimulation with ATP (Fig. 5C). Just like with GABA signaling, *Mtb* infection significantly reduced Ca^2+^ mobilization in scramble hBMDMs but further amplified the responses in GID/CTLH knockdown macrophages (Fig. 5C). Together, these functional data corroborated our host dual RNA-seq data sets which showed a specific enrichment of GABAergic and Ca^2+^ signaling pathways in *Mtb* infected GID/CTLH knockdown macrophages (Fig. 4C, Table S5).

**Fig.5.**
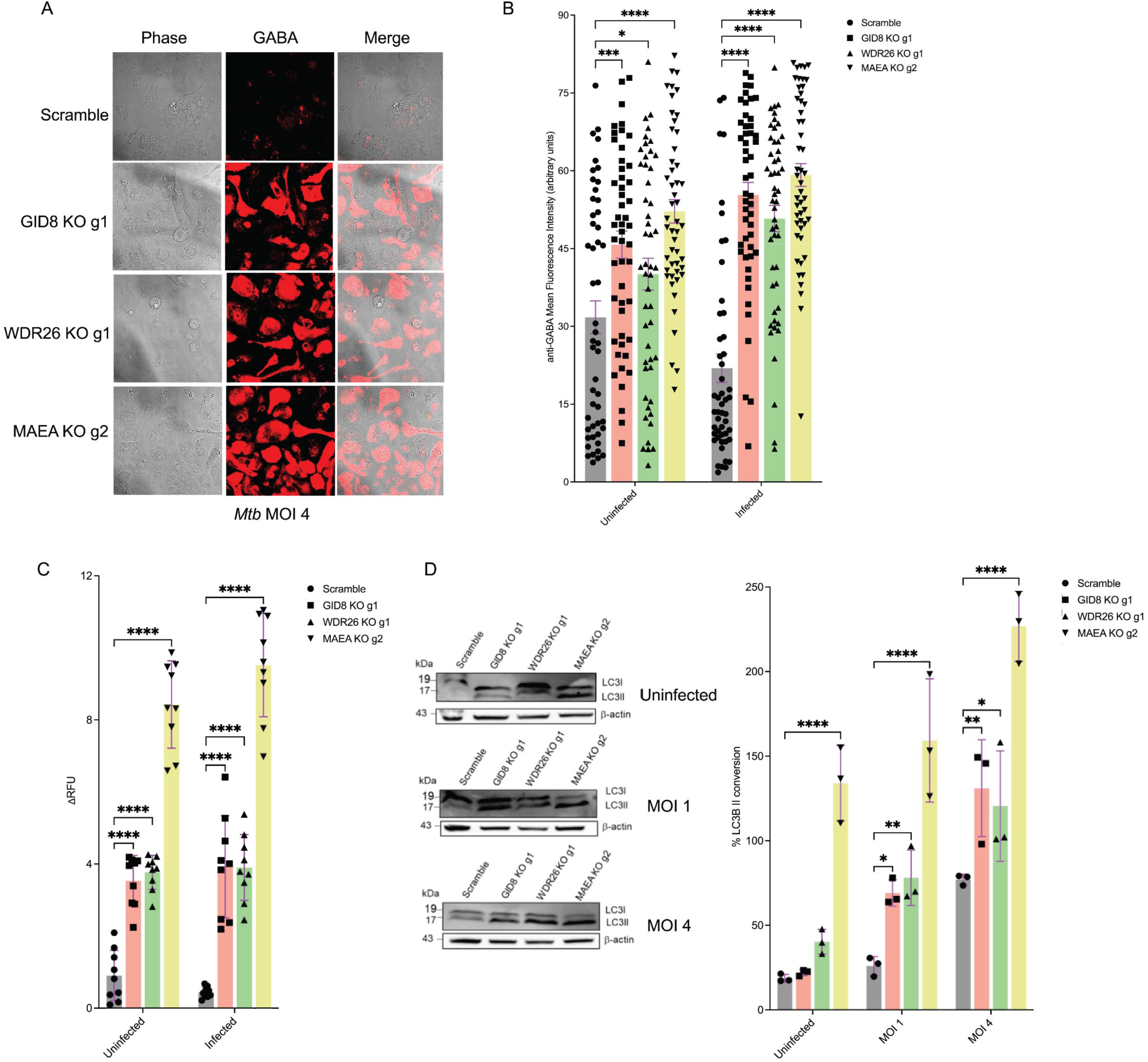
Increased autophagy, intracellular Ca^2+^ mobilization and GABAergic signaling in *Mtb* infected GID/CTLH knockdown macrophages. (A) Scramble or *GID8*, *MAEA* and *WDR26* knockdown hBMDMs were infected with the *Mtb Erdman hsp60::GFP* strain at MOI 4 for 24 hours. Fixed and permeabilized cells were then stained with anti-GABA (red) for confocal imaging. (B) Quantification of mean fluorescence intensities (MFI) of GABA staining in A and Fig. S7A. MFI was quantified in at least 50 cells for both scramble and knockdowns; n=50. *P < 0.05; ***P < 0.001; ****P < 0.0001. (C) Intracellular calcium mobilization in scramble or *GID8*, *MAEA* and *WDR26* knockdown hBMDMs uninfected or infected with *Mtb Erdman* strain at MOI 1 for 24 hours. hBMDMs were loaded with the fluorogenic calcium binding FluoForte^®^ dye and stimulated with adenosine triphosphate (ATP). Fluorescence was monitored for 5 minutes using a plate reader from which the delta relative fluorescence unit (ΔRFU) was obtained as described in methods; n=3 (3 technical replicates per repeat). ****P < 0.0001. (D) Western blot analysis of autophagic LC3I to LC3II conversion in uninfected and *Mtb* infected scramble or *GID8*, *MAEA* and *WDR26* knockdown hBMDMs at indicated MOIs (E) Densitometry quantification (in %) of LC3I to LC3II conversion in D and Fig. S7B, S7C as normalized to β-actin; n=3. *P < 0.05; **P < 0.01; ****P < 0.0001.

Increased GABA signaling in macrophages enhances antimicrobial responses by promoting autophagy.^55^ In fact, GABARAPL1, one of the upregulated GABA type A receptor in GID/CTLH knockdown hBMDMs (Table S4) is a homologue of the autophagy related protein 8 (*ATG8*).^71^ Genes involved in the autophagic process (*ATG9B, TRIM2, TRIM7 and TRIM16)* were also among the most significantly upregulated in *Mtb* infected GID/CTLH knockdown hBMDMs (Table S4). Moreover, in both uninfected and *Mtb* infected conditions, GID/CTLH knockdown hBMDMs display significantly increased AMPK phosphorylation (Fig. 3E). AMPK activates catabolic processes like autophagy as part of the energy stress response in cell starving conditions.^48^ These observations suggested that GID/CTLH knockdown hBMDMs could be more autophagic as part of their antimicrobial responses. We therefore checked autophagic flux in GID/CTLH knockdown hBMDMs by 1) immunoblotting to monitor the conversion of the microtubule-associated protein light chain 3 (LC3) from its LC3I to LC3II version 2) confocal imaging of anti-LC3 stained cells to quantify total LC3 puncta per individual cell and LC3 colocalizing with fluorescent *Mtb*. Indeed, we observed significantly increased autophagic flux in *Mtb* infected GID/CTLH hBMDMs as evidenced by high LC3I to LC3II conversion ratios when compared to scramble hBMDMs (Fig. 5D, Fig. S7B, S7C). The magnitude of LC3I to LC3II conversion proportionally increased based on the infecting bacteria MOI (Fig. 5D, Fig. S7B, S7C). Some GID/CTLH knockdown hBMDMs (*MAEA*) displayed increased LC3I to LC3II conversion even in uninfected conditions (Fig. 5D, Fig. S7B, S7C). Furthermore, *Mtb* infected GID/CTLH knockdown hBMDMs displayed significantly higher numbers of total LC3 puncta per cell as compared to scramble when we analyzed the anti-LC3 stained cells by confocal microscopy (Fig. S7D, S7E). More importantly, GFP expressing *Mtb* significantly colocalized with LC3 in GID/CTLH knockdown hBMDMs (Fig. S7D, S7F) when compared to scramble which indicated increased autophagic targeting of *Mtb* in these mutant macrophages. These data provided more functional proof that autophagy is indeed one of the main drivers of *Mtb* restriction in GID/CTLH knockdown hBMDMs. Given that increased GABA signaling can dampen the expression of proinflammatory markers in *Mtb* infected BMDMs,^55^ we also checked the expression of IL-1β and type 1 interferons (IFNB) in GID/CTLH knockdown macrophages during the early stages of infection. We already observed a significant downregulation of proinflammatory markers including Tnf and IL-1β in GID/CTLH knockdown macrophages in our RNA seq data which was collected 4 days post infection (Fig. S6). We thus concentrated our further analysis of the expression of IFNB and IL-1B in the first 24 hours of infection. qPCR analysis showed that *Mtb* infection caused a significant upregulation of IFNB mRNA in scramble macrophages 4 hours post infection, but this response diminished 24 hours post infection (Fig. S8A). On the contrary, IFNB mRNA expression was completely inhibited in GID/CTLH knockdown macrophages at both time points, and this was further confirmed by ELISA quantification of IFNB in cell supernatants 24 hours post infection (Fig. S8A). On the contrary, mRNA levels of IL-1β significantly increased in GID/CTLH knockdown macrophages during the early 4 hours of infection, but this quickly resolved after 24 hours of infection to non-significant levels (Fig. S8B). This contrasted with depleted levels of IL-1B mRNA 4 days post infection in our RNA seq data (Fig. S6). We were, however, able to confirm by ELISA that the resolution of IL-1β mRNA expression at 24 hours already corresponds with significantly lower levels of released IL-1β in culture supernatants as compared to scramble (Fig. S8B). Both the pro and mature version of IL-1β were equally affected when we checked each of the fractions by western blot (Fig. S8C) suggesting that the anti-IL-1β properties of GID/CTLH knockdown could be occurring upstream of the IL-1β caspase mediated processing. Taken together, these data provided more evidence that GID/CTLH knockdown impairs production of proinflammatory markers in *Mtb* infected macrophages.

### Concluding remarks

In this report, we have used a high throughput CRISPR genetic screen in primary macrophages to identify host effectors which, upon perturbation, improve the control of intracellular replication of *Mtb*. In the screen, pathways which are known to interfere with *Mtb* growth when perturbed such as mTOR signaling and the mitochondrial OXPHOs were specifically enriched. However, we also identified the mammalian GID/CTLH complex whose function in macrophage antimicrobial properties was previously unknown, to inversely correlate with intracellular *Mtb* replication upon knockdown.

The GID/CTLH complex is an evolutionary conserved member of the E3 ubiquitin proteasome system which was initially identified in yeast as a regulator of gluconeogenesis^34^ but is more functionally diverse in higher eukaryotes.^36^ With up to 10 functional members of the complex identified in mice and humans,^32^ the catalytic core of the complex is formed by the 2 RING proteins MAEA and RMND2A while GID8 (TWA1) is a predicted scaffold that holds the complex together.^32^ The entire complex has not been extensively studied in mice or humans. However, it is ubiquitously expressed in most mammalian cells^32^ which suggests its activities may have evolved to a rather greater complexity. As we show in this study, individual knockdown of up to 50% of the GID/CTLH complex members in murine and human macrophages renders these phagocytes hostile to the intracellular replication of both *Mtb* and *S. typhimurium*. It has been suggested that the GID/CTLH complex is involved in regulating cell proliferation, death, and survival pathways.^36^ SiRNA mediated knockdown of the GID/CTLH complex member RANBP9 made the cells more resistant to programmed cell death.^72,73^ Indeed, knockdown of the GID/CTLH complex in macrophages also rendered these phagocytes more resistant to *Mtb* induced necrotic cell death, a favorable host outcome which can limit bacterial spread and virulence.^10,13,14^ Moreover, GID/CTLH knockdown in macrophages reprogrammed these cells to a more glycolytic state with increased activation of AMPK in line with previous observations.^43,44^ Our results also identified additional anti-microbial responses upregulated in GID/CTLH complex knockdown macrophages, such as the NADPH oxidase, autophagy, GPCR signaling, iron efflux, GABAergic signaling and intracellular Ca^2+^ mobilization. GABAergic signaling has only recently been implicated in macrophage control of intracellular bacterial pathogens by promoting autophagy and inhibiting production of proinflammatory cytokines.^55^ Our results independently corroborate these findings as we observe increased GABA levels in GID/CTLH knockdown hBMDMs, which is similarly accompanied by enhanced autophagy and elevated anti-inflammatory properties. Dual RNA-seq also revealed that *Mtb* residing in GID/CTLH knockdown macrophages displayed transcriptional signatures consistent with nutritional stress as characterized by the upregulation of iron scavenger pathways, cholesterol breakdown, amino acid metabolism, fatty acid import and oxidation.

There is an increased interest in the incorporation of HDTs into anti-TB drug regimens as a possible means of shortening the duration of treatment.^30,53,74,75^ In theory, HDTs have the potential to 1) enhance host immune responses 2) promote macrophage control of intracellular *Mtb* 3) reduce non-productive and tissue injurious inflammation to improve lung function and 4) minimize host stresses that lead to induction of *Mtb* drug tolerance. ^75^ As we report, knockdown of the GID/CTLH complex in macrophages induces a broad range of anti-TB responses that suggest it would make an excellent target for HDTs against *Mtb*. Several members of this complex are upregulated in a variety of cancers and targeting this complex for cancer therapeutics is an area of current interest.^36,76–78^ As efforts continue to identify novel HDTs against TB,^1,53,75,79^ the GID/CTLH complex offers an attractive target and discovery efforts could benefit from the emerging therapeutic pursuit of the enzyme complex in the cancer field.

## Materials and methods

### Bacterial strains

The parent *Mtb Erdman strain* (ATCC 35801) was used in all CFU experiments. Fluorescent *Mtb* reporters in the *Erdman* background; *Mtb Erdman smyc’::mCherry* and *Mtb Erdman hsp60::GFP* have been described previously.^80,81^ The *Mtb Erdman*-Lux strain was generated by transformation with the pMV306G13+Lux plasmid.^82^ *Mtb* was grown to log phase at 37°C in MiddleBrook 7H9 broth supplemented with 10% oleic acid/albumin/dextrose/catalase (OADC Enrichment; Becton, Dickinson and Company), 0.2% glycerol, and 0.05% tyloxapol (Sigma-Aldrich). Reporter strains were maintained in the presence of an appropriate antibiotic; *Mtb Erdman*-Lux *Mtb* strain: 25μg/ml kanamycin, *Mtb Erdman hsp60::GFP*: 25μg/ml kanamycin and 50μg/ml hygromycin, *Mtb Erdman smyc’::mCherry*: 50μg/ml hygromycin. For *Salmonella* experiments, wild type *S.* Typhimurium (CA32; ATCC 14028s) and the BFP expression strain *S.*Typhimurium CA4705 phoN::BFP^41^ were used. *Salmonella* strains were propagated at 37°C in Luria-Bertani (LB) broth.

### Mammalian cells and culture

Murine Hoxb8-Estradiol (ER) responsive Cas9^+^ eGFP conditionally immortalized myeloid progenitor cells were generated as previously described.^22^ Hoxb8-ER cells were maintained in RPMI (Corning^®^) supplemented with 10% fetal bovine serum (FBS), 2 mM L-glutamine, 1 mM sodium pyruvate, 20ng/ml murine GM-CSF (PeproTech), 0.5 μM β-estradiol, 10mM HEPEs and 1% penicillin/streptomycin. To obtain hBMDMs from Hoxb8-ER myeloid progenitors, cells were rinsed twice with 1x PBS to completely remove β-estradiol and resuspended in BMDM differentiation media; DMEM (Corning^®^) supplemented with 10% FBS, 15% L-cell conditioned media, 2 mM L-glutamine, 1 mM sodium pyruvate, and 1% penicillin/streptomycin at 37°C for 6-7 days. Cells were maintained at a density of ∼0.5 x 10^6^ cells/ml during differentiation. The HEK-293FT cell line was purchased from Invitrogen and cultured in DMEM supplemented with 10% FBS, 1% non-essential amino acids, 2 mM L-glutamine, 1 mM sodium pyruvate, and 1% penicillin/streptomycin. Human monocytes were commercially obtained from the University of Nebraska Medical Center Elutriation Core and differentiated to HMDMs in DMEM medium with 2mM L-glutamine, 1 mM sodium pyruvate, 10 mM HEPES, 1% penicillin/streptomycin and 10% pooled human serum (Sera Care).

### Infection of macrophages with *Mtb*

hBMDMs were infected with *Mtb* as previously described.^83^ Briefly, log phase bacteria were pelleted, resuspended in basal uptake buffer (25 mM dextrose, 0.5% bovine serum albumin, 0.1% gelatin, 1 mM CaCl2, 0.5 mM MgCl2 in PBS) and syringed with a tuberculin syringe for at least 20 times. Bacteria were then resuspended in antibiotic free macrophage media before infection at specified MOIs.

### CFU quantification of intracellular bacteria

Confluent macrophage monolayers (human and murine) were infected with *Mtb* at an appropriate MOI. After 3-4 hours, extracellular bacteria were removed by washing with fresh macrophage media at least 3 times. At indicated time points, macrophages were lysed with 0.01% SDS in water for 15 minutes and serially diluted in 0.05% Tween-80 in 1x PBS. Lysates were plated on 7H10 OADC agar plates and incubated at 37^0^C for CFU counting 3-4 weeks later.

### Cloning of sgRNAs, lentiviral production and transduction

sgRNAs for CRISPR mediated gene knockdowns (Table S3) were selected from the Brie murine knockout library^23^ or ordered in predesigned form from IDT for human CRISPR RNP targets. For lentiviral delivery of sgRNAs, oligo pairs were annealed and cloned into the pLentiGuide-Puro (Addgene #52963). sgRNA inserts were confirmed by sequencing. To generate lentivirus particles for transduction, pLentiGuide-Puro plasmids with sgRNAs of interest were co-transfected with Invitrogen packaging plasmids (pLP1, pLP2, and pLP/VSVG) in 293FT cells using the Lipofectamine 3000 lentiviral production protocol according to manufacturer’s guidelines. To transduce Hoxb8 Cas9^+^ eGFP progenitors, 2.5 x 10^5^ cells in 12 well plates were spininfected at 1000xg for 90 minutes at 32°C in the presence of 10 µg/ml protamine sulfate. 48 hours post transduction, 8 µg/ml puromycin was added and maintained for 4 days. After the 4-day selection period, puromycin resistant cells were briefly expanded for 1-2 days and frozen for future experiments. To quantify CRISPR editing efficiencies, total genomic DNA was prepared from cell pellets using the Qiagen DNeasy Blood and Tissue kit from which genomic sites surrounding the sgRNA target sites were PCR amplified using the Invitrogen Platinum *Taq* polymerase. Population level editing efficiencies were estimated using ICE^39^ and western blot analysis of differentiated hBMDMs.

### CRISPR screen knockout library preparation

The murine Brie knockout library was obtained from Addgene (#73633).^23^ The library (in pLentiGuide-Puro background) contains 78,637 sgRNAs targeting 19,674 genes (4 sgRNAs/gene) with an additional 1000 non-targeting sgRNA controls. The library was amplified using the Moffat’s lab CRISPR knockout library amplification protocol.^84^ To generate lentivirus, 293FT cells were transfected as described above, scaling up tissue culture flasks depending on the amount of virus required to achieve desired library coverage. ∼180 million Cas9 eGFP Hoxb8 progenitor cells were then transduced with the lentiviral library in the presence of 8 µg/ml polybrene at an MOI of ∼0.3 (8 12-well plates, 2 x 10^6^ cells/well, >700x library coverage) to ensure that only a single gene was targeted in every cell. 48 hours post-transduction, cells were selected with 8 µg/ml puromycin for 4 days and frozen in multiple aliquots of >40 million cells (>500x library coverage). Before screening, each aliquot was thawed and allowed to recover in Hoxb8 media for 3-4 days. Cells were then rinsed in 1x PBS at least 2 times before transfer into BMDM differentiation media to generate hBMDMs knockout libraries.

### Mtb infection of hBMDM knockout library, antibody staining and flow sorting

The hBMDM knockout library was infected with the *Mtb Erdman* strain at MOI 1 for 4 days. The screen was carried out in 3 independent replicates with ∼200 million cells per replicate (2 T-300 flasks). On day 4, all the cells were harvested by initially pretreating cell monolayers with Accutase for 5-10 minutes at room temperature followed by scrapping in cold 1x PBS. Harvested cells were stained with the live/dead viability dye eFlour 506 (Invitrogen; 1:1000) and the PE anti-mouse/human CD11b antibody (Biolegend; 1:500) for 30 minutes at 4^0^C. Stained cells were fixed in paraformaldehyde (PFA) for > 1 hour before flow cytometry activated sorting (FACS) of cells in the live gate. FACS was carried out on a Sony MA900 sorter and a minimum of 10 million cells were sorted per replicate to achieve a >150x library coverage.

### DNA extraction, barcode amplification, sequencing and analysis

Genomic DNA was extracted from the sorted cells as well as the unperturbed input library at full coverage (>300x library coverage) using a modified salt precipitation protocol as described previously.^85^ Briefly, for 3 x 10^7^-5 x 10^7^ frozen cell pellets (scaled down proportionally for lower cell numbers), 6 ml of NK Lysis Buffer (50 mM Tris, 50 mM EDTA, 1% SDS, pH 8) and 30 μl of 20 mg/ml Proteinase K (Qiagen) were added to the cells in a 15 ml falcon tube and incubated at 55^0^C overnight to lyse the cells and de-crosslink PFA fixation. Next day, 30 μl of 10 mg/ml RNAse A diluted in NK lysis buffer was added to the lysed sample and incubated at 37^0^C for 30 minutes. Samples were chilled on ice before addition of 7.5 mM ice cold ammonium acetate to precipitate proteins. After a brief vortex, insoluble protein fractions were pelleted by centrifugation at ≥ 4,000 x *g* for 10 minutes. The supernatant was transferred to a new 15ml falcon tube and DNA was precipitated by Isopropanol. Precipitated DNA was washed 3 times with 70% ethanol and airdried. Dried pellets were dissolved in sterile nuclease free water and DNA concentration was measured on a Nanodrop 1000 (Thermo Scientific). Amplification of sgRNAs cassettes from the extracted DNA was performed using Illumina compatible primers from IDT as described previously^23^ with some minor modifications. PCR amplification was carried out using KAPA HIFI Hotstart PCR (Roche) with the following reaction mixture in a 50 μl volume: 10 μL 5x reaction buffer, 1.5 μL dNTP, 1.5 μL P5 primer mix, 10 μM, 1.5 μL of P7 primer 10 μM, 2.5 ul DMSO, 0.5 μL polymerase, up to 2 μg of genomic DNA or 10 ng of plasmid DNA, up to 50 μL with water. PCR cycling conditions were as per manufactures protocol. Target PCR products were gel extracted using the Qiagen gel extraction kit and re-purified with the GeneJET PCR Purification kit (Thermo) before sequencing on an Illumina NextSeq500. Fastq files were mapped to the sgRNA library Index using MAGeCK-VISPR which allows for automated trimming of adaptors and identification of sgRNA length.^24–26^ Ranked sgRNA and gene hits were similarly obtained with the MAGeCK-VISPR workflow using Robust Rank Aggregation.

### GO term enrichment

GO term enrichment was carried out in R using the enrichGO function of clusterProfiler.^51^ Enriched BP, CC or MF were filtered based on an adjusted p value of < 0.05. In some cases, top enriched terms were visualized by tree plots of related GO clusters

### Seahorse flux analyses

Extracellular flux analyses to measure oxygen consumption rates (OCRs) and extracellular acidification rates (ECARs) were performed using the Seahorse-8 XFp flux analyzer (Agilent). hBMDMs were plated at a density of 1 x 10^5^ cells/well in 8-well Seahorse plates overnight. Cells were then infected with *Mtb Erdman* strain at MOI 1 or left uninfected for another 24 hours. 1 hour before the assay, hBMDM media was replaced with Seahorse base medium (without phenol red and sodium bicarbonate, with 5 mM HEPES) and placed at 37°C in a non-CO2 incubator. For the Mito stress test to measure OCR, 10mM glucose, 2mM L-glutamine and 1mM sodium pyruvate were added to the Seahorse base medium while only glutamine was added for the Glycolysis stress test to measure ECARs. OCR and ECAR measurements were performed in line with specific kit instructions by a sequential injection of oligomycin (2.5μM), FCCP, fluoro-carbonyl cyanide phenylhydrazone (1.5μM), rotenone/antimycin A (0.5μM) for the Mito stress assay and glucose (10mM), oligomycin (2.5μM), 2-Deoxy-D-glucose (50mM) for the Glycolysis stress assay. 3 measurements were taken under basal conditions and after each drug injection.

### Western blot analysis

Cells were washed in ice cold 1x PBS and lysed with RIPA buffer (Thermo Fisher) containing protease inhibitors (Roche) for 30 minutes at 4^0^C with gentle agitation. The Pierce™ Rapid Gold BCA Protein Assay Kit (Thermo Fisher) was used to measure the concentration of protein lysates. Equal amounts of protein (30μg) were separated by SDS PAGE on a 4-20% gradient gel and transferred to a nitrocellulose membrane. The membrane was blocked for non-specific binding by incubating with 5% non-fat milk in PBST (0.1% tween in 1x PBS) for at least 30 minutes. Blots were washed and incubated with primary antibodies diluted in 2% non-fat milk in PBST at 4^0^C overnight. The following primary antibodies were used in western blot analyses. Anti-AMPKα Mab (1:1000, Cell signalling Technology), anti-Phospho-AMPKα Mab (1:500, Cell signalling Technology), anti-β Actin (1:1000, Cell Signalling Technology), anti-LC3B (1:1000, Cell Signalling Technology), anti-GID8 (1:1000, Proteintech), anti-YPEL5 (1:1000, Proteintech), anti-WDR26 (1:300, Bioss), anti-UBE2H (1:1000, Proteintech), anti-MAEA (1 μg/ml, R&D Systems), anti-IL-1β (1:1000, Cell Signalling Technology). Secondary antibodies used were horseradish peroxidase conjugated polyclonal donkey anti-sheep IgG (1:1000, R&D Systems) and anti-rabbit/mouse StarBright Blue 700 (1:2500, Biorad). Blots were developed by the SuperSignal™ West Atto Ultimate Sensitivity Substrate (Thermo Fisher) or directly imaged in case of fluorescent secondary antibodies. Images were acquired on a ChemiDoc MP imaging system (Biorad). Western blot band intensities were quantified in ImageJ.

### Dual RNA-sequencing of hBMDM Mtb infected macrophages

hBMDMs were infected with the *Mtb Erdman smyc’::mCherry* reporter strain at MOI 0.5. 4 days post infection, cells were harvested and stained with the live/dead viability dye eFlour 506 (Invitrogen; 1:1000) and the PE anti-mouse/human CD11b antibody (Biolegend; 1:500) for 30 minutes at 4^0^C. Live mCherry Cd11b positive cells were sorted on a Sony MA900 sorter. 250, 000 *Mtb* infected sorted macrophages were spun down, resuspended in 150 μl sorter buffer and transferred to a tube containing 600 μl Trizol. Samples were mixed and incubated at room temperature for 5 minutes followed by centrifugation at ∼12 000g for 20 minutes to pellet the bacteria. 600 μl of Trizol supernatant (host RNA) was then transferred to a new tube. Fresh 400 μl of Trizol was added to the tube containing the bacterial pellet together with 150 μl of zirconia beads. Bead beating was performed on a FastPrep-24™ bead beater (MP biomedicals) in 2 cycles of 1 minute resting 2 minutes on ice between each cycle. 40% of host RNA (∼240 μl) was then added back to the original bead containing tube which allows for a good proportion capture of both host and bacteria transcripts by getting rid of some of the host RNA.^49^ Chloroform was added (200 µL of chloroform for 1 mL of Trizol) to the tubes which were mixed thoroughly and spun at 12, 000g for 15 minutes at 4^0^C. Total RNA was purified from the aqueous phase using the Zymo RNA clean and concentrator-5 kit according to manufactures instructions. Ribosomal RNA (rRNA) depletion in the RNA samples was performed using the QIAseq FastSelect pan-bacterial and H/M/R kits in the duo-depletion mode. Libraries were prepared from rRNA depleted samples using the NEBNext Ultra II Directional RNA Library Prep Kit (New England Biolabs) and sequenced on the Novaseq 6000 S4 (Illumina). Paired end sequenced reads were mapped to the mouse reference genome (GRCm39) and the *Mtb* H37Rv genome (ASM19595v2) using STAR-2.7.10b. Raw counts were obtained using HTseq with GRCm39 and ASM19595v2 annotations. Normalization of read counts and differential expression analysis were performed in R using the DESeq2 package. Pathway enrichment was performed on DE genes using clusterProfiler as described in the prior section.

### Intracellular Ca2+ mobilization assay

hBMDMs were seeded in 96-well black clear bottom plates at a density of 1 x 10^5^ cells/well and incubated at 37 °C in 5% CO2 overnight. Next day, macrophage media was discarded and replaced with the Fluoforte® dye (Enzo Life) in 100 μl Hank’s balanced salt solution (HBSS) in the presence of a dye efflux inhibitor. After 1 hour of incubation (45 minutes at 37 °C and 15 minutes at room temperature), 20 μl of 6 μM ATP was added to each well for a final concentration of 1 μM. Fluoforte® fluorescence (excitation 490 nM/emission 525 nM) was quantified at time 0 (baseline) and after 5 minutes on an Envision plate reader (PerkinElmer). Normalized relative fluorescence units (ΔRFU) were calculated for each well using the following formula:

ΔRFU = (Fluorescence taken after 5 minutes of ATP stimulation-Baseline fluorescence taken at the start of the assay)/ Baseline fluorescence taken at the start of the assay

### Immunofluorescence and confocal microscopy

Macrophage monolayers in Ibidi eight-well chambers were infected with *Mtb* at an appropriate MOI as described in prior sections. At specified time intervals, cells were fixed with 4% PFA for > 1 hour and permeabilized with 0.2% Triton X-100 (Sigma-Aldrich) for 15 minutes. Permeabilized cells were blocked with 3% Bovine Serum Albumin (BSA) for 1 hour and incubated with primary antibody diluted in 3% BSA at 4 °C overnight. Next day, cells were washed with 1x PBS 3 times and incubated with secondary antibodies for 1 hour at room temperature. Primary antibodies used were anti-GABA (1:400, Sigma-Aldrich) and anti-LC3B (1:1000, Cell Signalling Technology). The Alexa Fluor™ 647 goat anti-rabbit secondary antibody (1:500, Thermo Scientific) was used. After mounting, images were acquired using a Leica SP5 confocal microscope. Z-stacks were reconstructed in ImageJ from which mean fluorescence intensities (MFI) for individual cells or colocalizations were performed.

### CRISPR RNP delivery in human monocyte derived macrophages

Isolated human monocytes were left in culture for 24 hours before transfection. On the day of transfection, cells were harvested, washed with 1x PBS and resuspended in P3 Primary Cell Nucleofection Buffer (Lonza) at a concentration of 5 × 10^6^ cells in 20 μl for each electroporation. Equal volumes of 100 μM Alt-R crRNA and Alt-tracrRNA (IDT) were annealed to form sgRNA duplexes by incubation at 95°C for 5 minutes followed by cooling to room temperature for > 15 minutes. To form CRISPR RNPs, annealed sgRNA duplexes were mixed with IDT Alt-R S.p. Cas9 Nuclease V3 at a 3:1 molar ratio and incubated for at least 20 minutes at room temperature. For nucleofection, 4 μl of CRISPR RNP was mixed with the 20 μl cell suspension in P3 buffer and transferred into the supplied nucleofector cassette strip. The strip was inserted into the Lonza 4D-Nucleofector and nucleofected with the Buffer P3, CM-137 conditions as previously described.^42^ After nucleofection, cells were transferred into petri dishes and allowed to complete differentiation to HMDMs for 5-6 days. Knockdown efficiencies were confirmed in HMDMs by ICE analysis and western blot.

### Quantitative RT-PCR

Total RNA was isolated from uninfected or *Mtb* infected hBMDMs using Trizol alongside the Zymo RNA clean and concentrator-5 kit according to manufactures instructions. cDNA was synthesized from ∼ 1 μg of extracted RNA using an iScript cDNA synthesis kit (Bio-Rad). RT-PCR was performed for IFNB and IL-1β on the Applied Biosystems PRISM 7500 qPCR machine. Beta 2 microglobin was used as a reference housekeeping gene. Predesigned qPCR primers were purchased from IDT. PCR reactions were carried out using iTaq Universal SYBR Green Supermix (Bio-Rad).

### ELISA

hBMDMs in 96 well plates were infected with *Mtb* at an appropriate MOI for 24 hours. Cell culture supernatants were collected from infected cells and assayed for IFNB and IL-1β using the IFNB and IL-1β R&D ELISA kits according to manufactures instructions.

### General statistics

Basic statistical tests were carried out in GraphPad Prism 10. Comparisons between more than two groups were analyzed using one-way or two ANOVA alongside the Dunnett’s multiple comparison test. P values less than 0.05 were considered statistically significant.

## Acknowledgments

We thank Dr. Jen K. Grenier and Ann E. Tate from the Cornell BRC Transcriptional Regulation and Expression Facility for ongoing support to improve the Dual RNA-sequencing protocols. Some cartoons in the graphical abstract were obtained from Biorender.com. This work was supported by grants from the National Institutes of Health (AI155319, AI162598, and OD032135), Bill and Melinda Gates Foundation and the Mueller Health Foundation to D.G.R., and by a grant from the National Institutes of Health (AI172433) to C.A. EJ was supported by T32AI007349.

