## Supplemental Data for "Genome-wide screen of *Mycobacterium tuberculosis-*infected macrophages identified the GID/CTLH complex as a determinant of intracellular bacterial growth"

**Supplementary tables**

Table S1. Hits from *Mtb* induced cytotoxicity macrophage survival CRISPR screen.

Table S2. Enriched GO terms from CRISPR screen hits

Table S3. sgRNAs Primers and ICE scores for GID/CTLH targets in mice and human

Table S4. Differentially expressed genes in GID8, MAEA and WDR26 knockdown *Mtb* infected macrophages

Table S5. Enriched GO terms amongst commonly upregulated and downregulated genes in GID8 MAEA and WDR26 knockdown macrophages

Table S6. *Mtb* differentially expressed genes in GID8 MAEA and WDR26 knockdown infected macrophages

### Supplementary figures

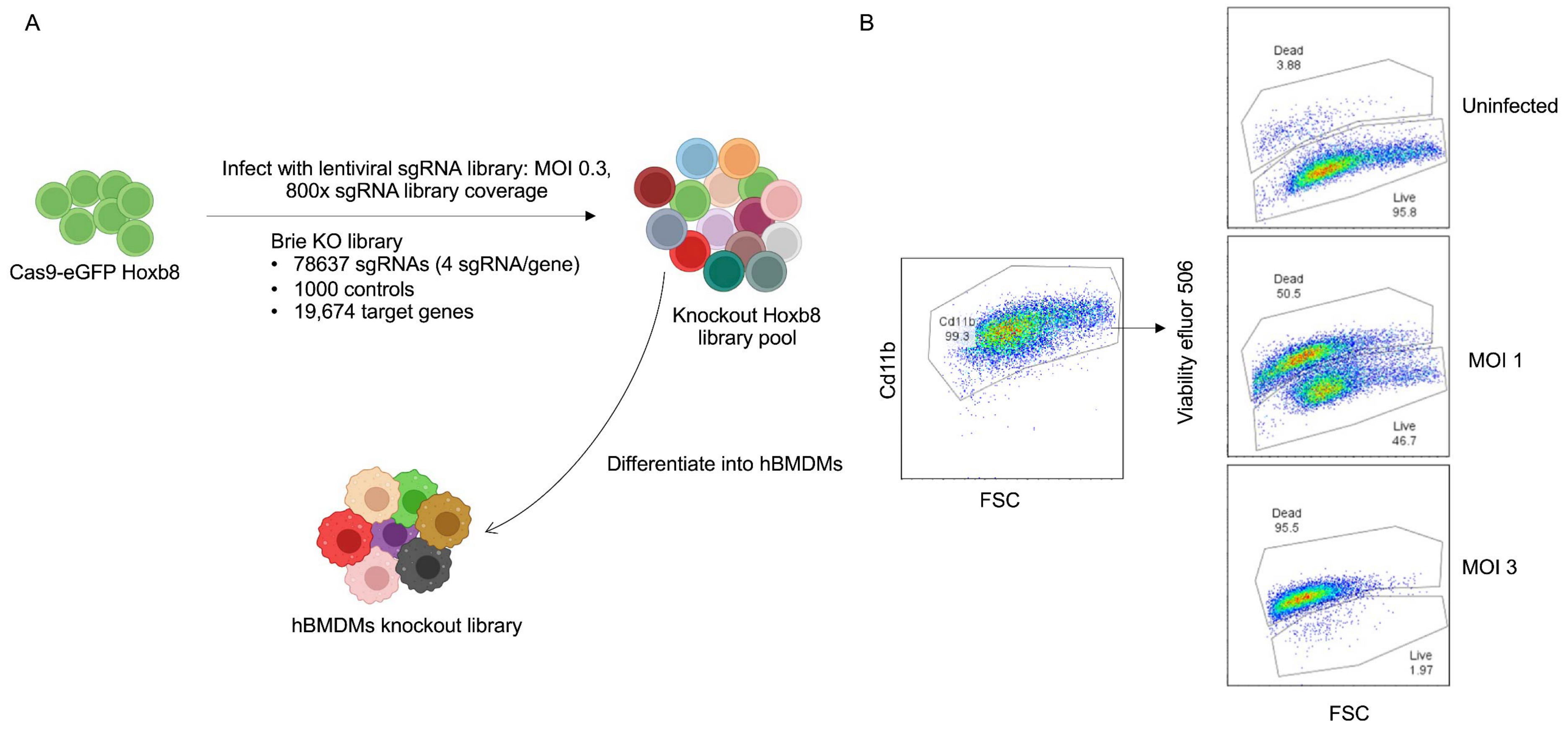

**Fig. S1. Related to Fig.1**

(A) Workflow for generation of Hoxb8 progenitor cells CRISPR knockout library. ~180 million Cas9 eGFP Hoxb8 progenitor cells were transduced with the Brie knockout library at an MOI of ~0.35 to achieve an ~800x library coverage. Cells were selected with puromycin for 4 days and frozen in aliquots of ~40 million cells (~500x library coverage). For screening, each aliquot was thawed and allowed to recover in Hoxb8 media for 3-4 days before transfer into hBMDM media to generate hBMDMs knockout libraries

(B) Quantification of *Mtb* induced cytotoxicity by flow cytometry. hBMDMs were infected as in Fig. 1A. All harvested cells were stained with the macrophage surface marker Cd11b and the live/dead eflour viability dye in 1x phosphate buffered saline (PBS) at 4 degrees for 30 minutes. Cells were then fixed in 4% paraformaldehyde (PFA) for at least 1 hour before analysis on a flow cytometer.

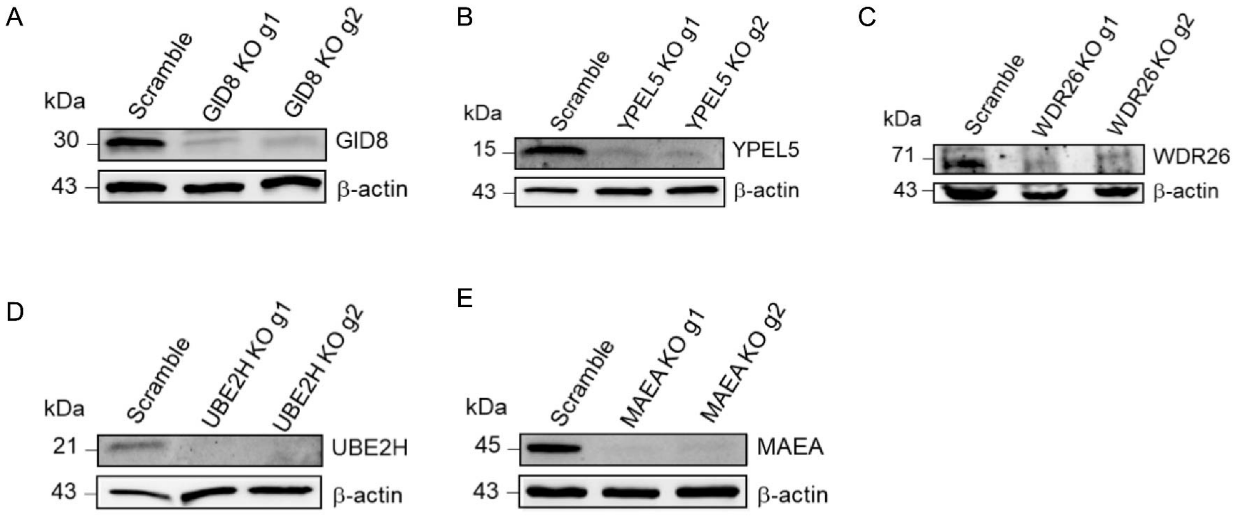


**Fig. S2. Related to Fig.2**

(A-E) Confirmation of target protein depletion in GID/CTLH knockdown hBMDMs derived from Hoxb8 parental lines (Table S3) by western blot. Analysis is for each of the 2 targeting sgRNAs compared to scramble, *GID8* (A), *YPEL5* (B), *WDR26* (C), *UBE2H* (D), *MAEA* (E).

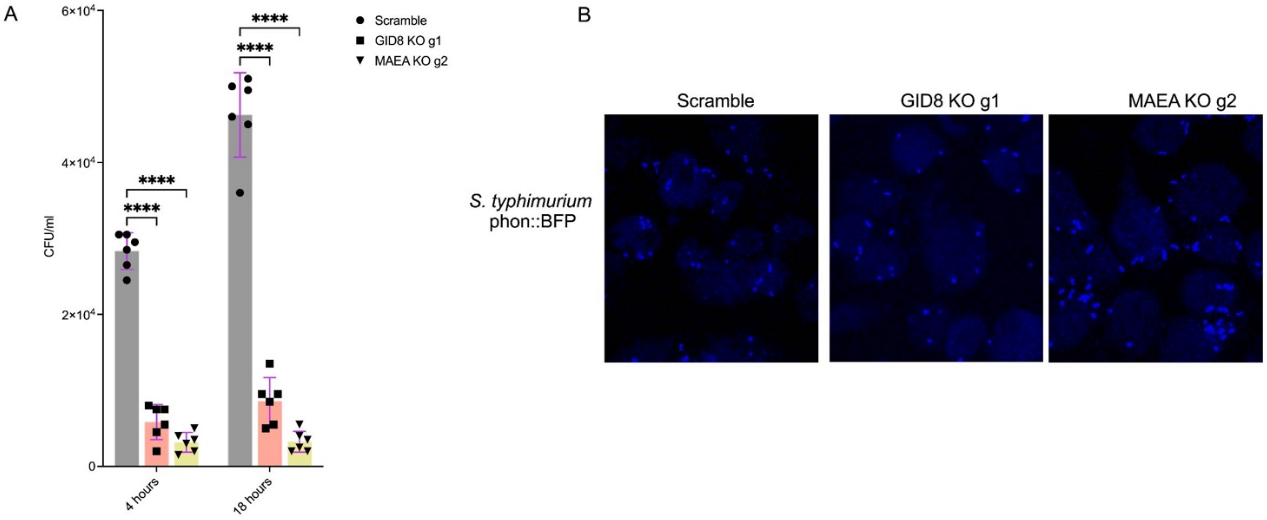


**Fig. S3. Related to Fig.2. Intracellular replication of *Salmonella typhimurium* in GID/CTLH knockdown macrophages**

(A) Intracellular growth of *S. typhimurium* in scramble or *GID8* and *MAEA* knockdown macrophages in the gentamicin protection assay. hBMDMs were infected with the *S. typhimurium* CA32 strain at MOI 10 for 30 minutes. Bacteria was recovered from lysed cells and plated for CFUs at the indicated time points; n =3 (2 technical replicates per repeat). ****$P < 0.0001$.

(B) Scramble or *GID8* and *MAEA* knockdown macrophages were infected with the *S. typhimurium* CA4705 phoN::BFP strain as in A. Cells were fixed 1 hour post infection and confocal images were acquired to visually estimate bacterial uptake efficiencies.

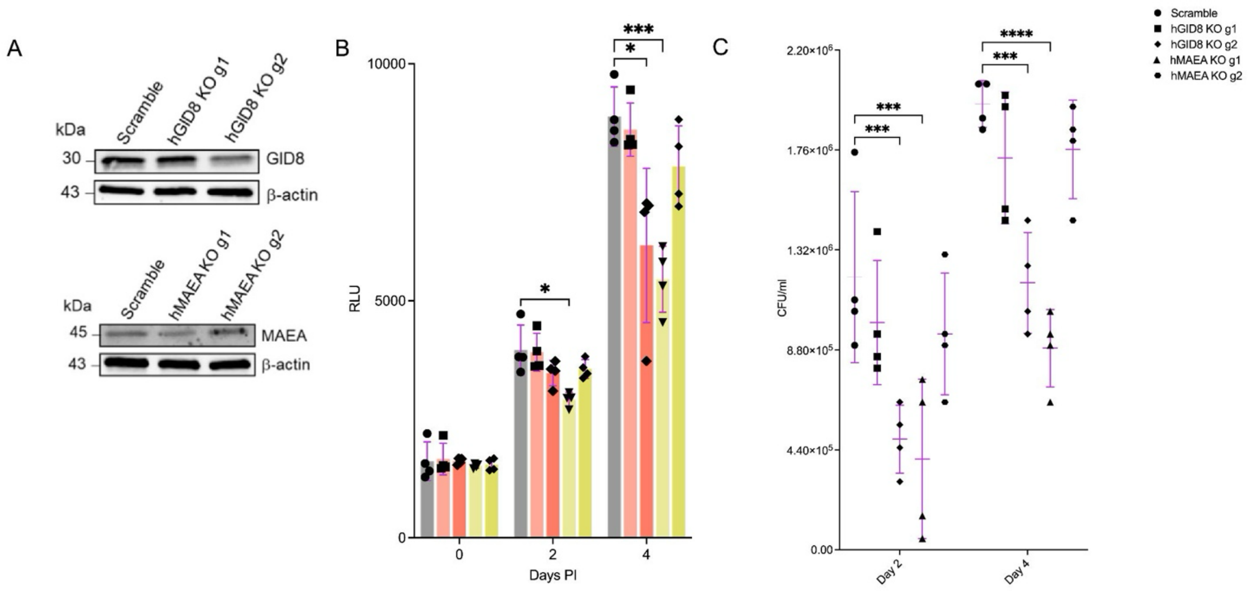


**Fig. S4. Related to Fig.2.** ***Mtb*** **growth restriction in human monocyte derived macrophages with** ***GID8*** **and** ***MAEA*** **gene knockdowns**

(A) Western blot analysis showing *GID8* and *MAEA* knockdown efficiencies by CRISPR RNPs in human monocyte derived macrophages (HMDMs)

(B) Quantification of *Mtb* replication in scramble or *GID8* and *MAEA* knockdown HMDMs using the *Mtb Erdman*-Lux strain. HMDMs were infected at MOI 0.5 and luciferase measurements were taken on the indicated days post infection; n=2 (2 technical replicates per repeat). *P < 0.05; ***P < 0.001.

(C) Scramble or *GID8* and *MAEA* knockdown HMDMs were infected with *Mtb Erdman* strain at MOI 0.4. CFUs were plated on day 2 and 4 to determine intracellular *Mtb* replication rates; n=2 (2 technical replicates per repeat). ***P < 0.001; ****P < 0.0001.

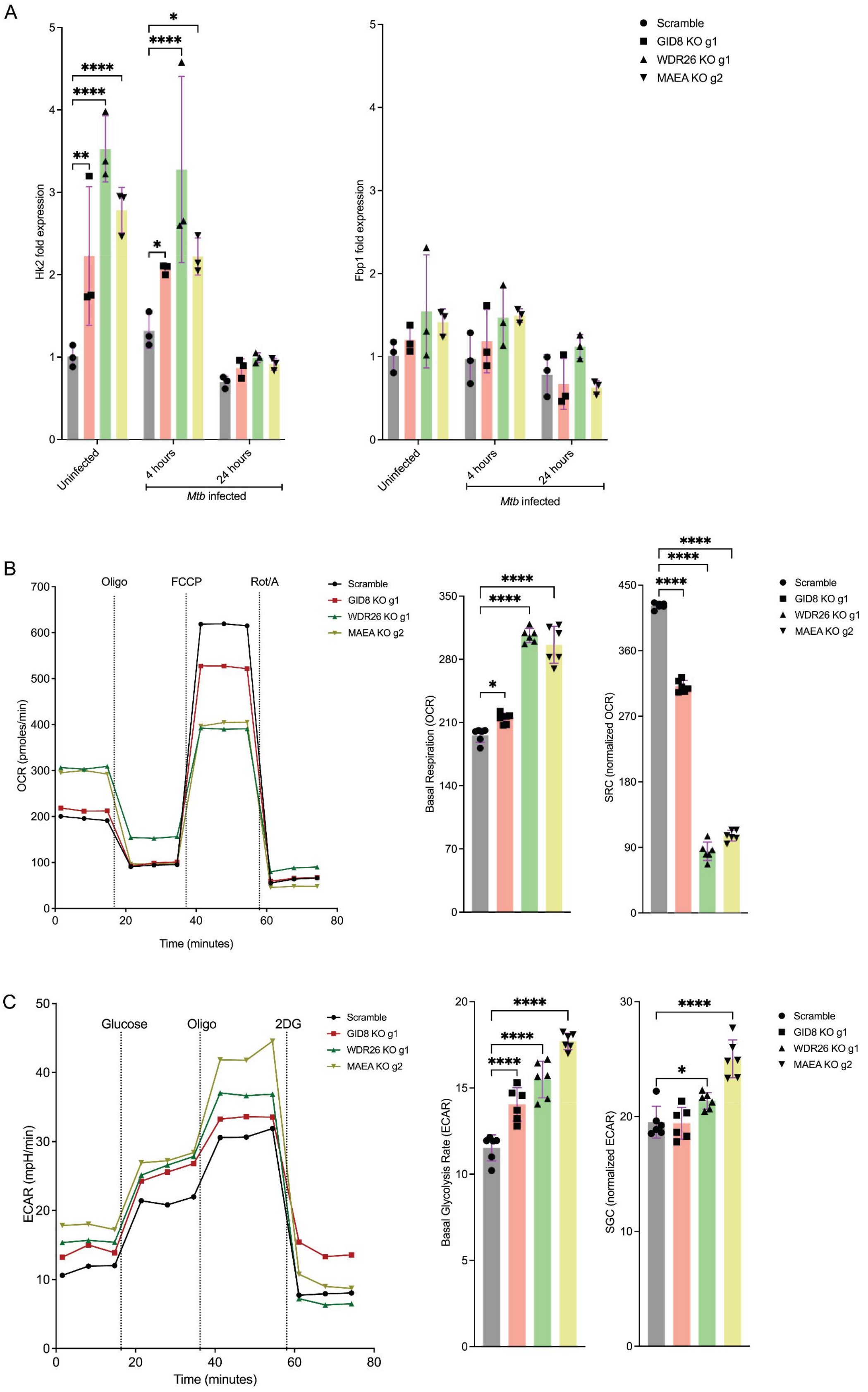
A
Hk2 fold expression
Fbp1 fold expression
Uninfected
4 hours
24 hours
Mtb infected
Scramble
GID8 KO g1
WDR26 KO g1
MAEA KO g2
B
Oligo
FCCP
Rot/A
OCR (pmoles/min)
Time (minutes)
Basal Respiration (OCR)
SRC (normalized OCR)
C
Glucose
Oligo
2DG
ECAR (mpH/min)
Time (minutes)
Basal Glycolysis Rate (ECAR)
SGC (normalized ECAR)

106

**Fig. S5. Related to Fig.3**

(A) qPCR analysis of hexokinase 2 (*HK2*) and fructose bisphosphate 1 (*FBP1*) mRNA expression in scramble or GID8, *MAEA* and *WDR26* knockdown macrophages. RNA was extracted from hBMDMs uninfected or infected with the *Mtb Erdman* strain at MOI 4 for 4 or 24 hours; n=3. *P < 0.05; **P < 0.01; ****P < 0.0001.

(B) Seahorse flux analyses in uninfected scramble or *GID8*, *MAEA* and *WDR26* knockdown macrophages using the Cell Mito Stress Kit as in Fig. 3A, 3B

(C) Seahorse flux analyses in uninfected scramble or *GID8*, *MAEA* and *WDR26* knockdown macrophages using the Glycolysis Stress Test kit as in Fig. 3C, 3D

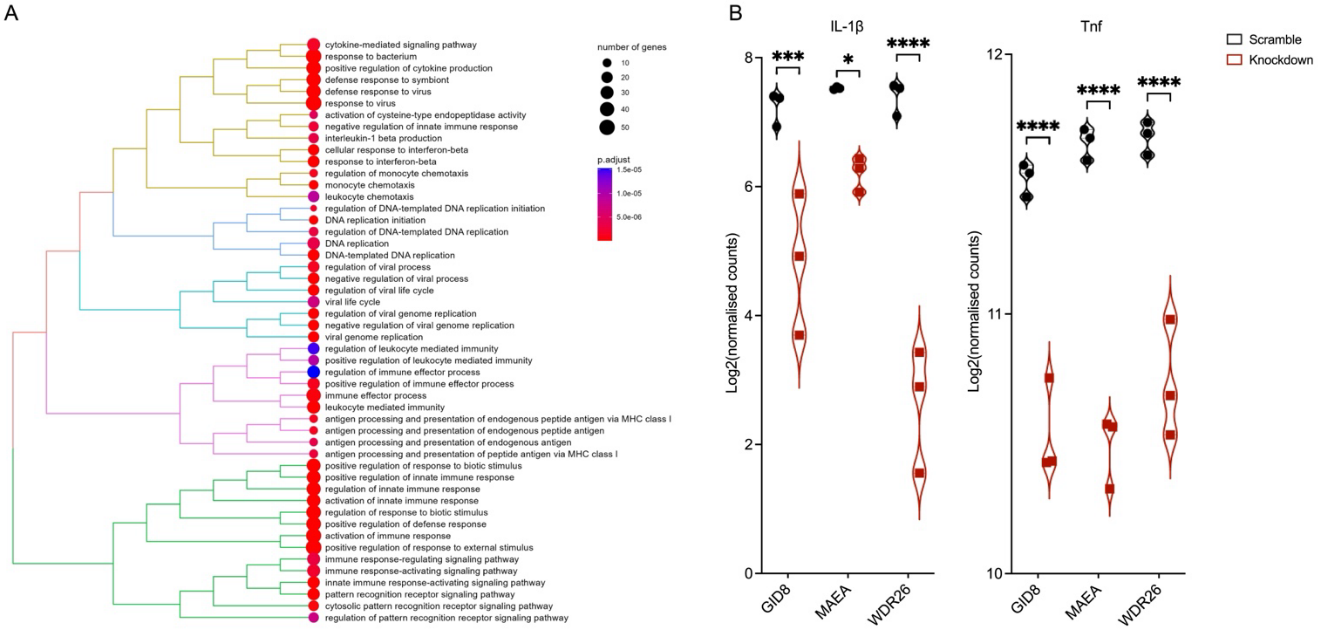


**Fig. S6. Related to Fig.4**

(A) Tree plot of top 50 functionally enriched gene ontology terms (biological process) amongst the 421 commonly downregulated genes in *GID8*, *MAEA* and *WDR26* knockdown hBMDMs

(B) Violin plot showing expression (log2 normalized counts) of interleukin 1b (IL-1β) and tumor necrosis factor (Tnf) in *Mtb* infected *GID8*, *MAEA* and *WDR26* knockdown macrophages 4 days post infection; n=3. *P < 0.05; ***P < 0.001; ****P < 0.0001.

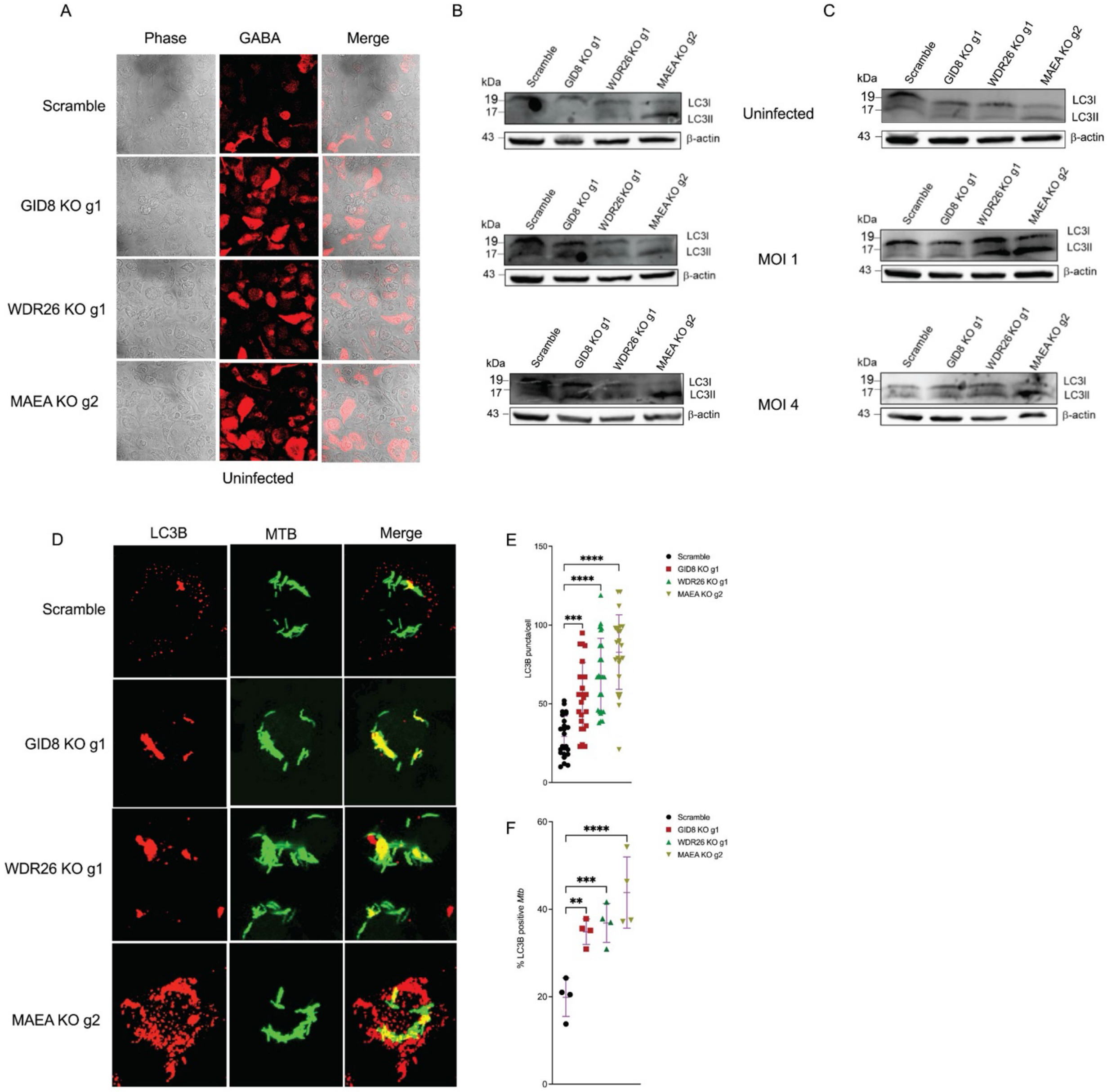


**Fig. S7. Related to Fig.5**

(A) Uninfected scramble or *GID8*, *MAEA* and *WDR26* knockdown hBMDMs were fixed and stained with anti-GABA (red) as in Fig. 5A

(B, C) Western blot images of the biological replicates for the LC3I to LC3II conversion analysis as in Fig. 5D

(D) Scramble or *GID8*, *MAEA* and *WDR26* knockdown hBMDMs were infected with *Mtb Erdman hsp60::GFP* strain at MOI 4 for 24 hours. Fixed and permeabilized cells were stained with anti-LC3B (red) for confocal imaging

(E) Quantification of LC3B punta per individual cell for confocal images in D. At least 24 cells were counted for both scramble and GID/CTLH hBMDMS knockdowns; n=24. ***P < 0.001; ****P < 0.0001

(F) Percentage GFP expressing *Mtb* colocalizing with LC3B for confocal images acquired as in D in at least 4 different fields; n=4. **P < 0.01; ***P < 0.001; ****P < 0.0001.

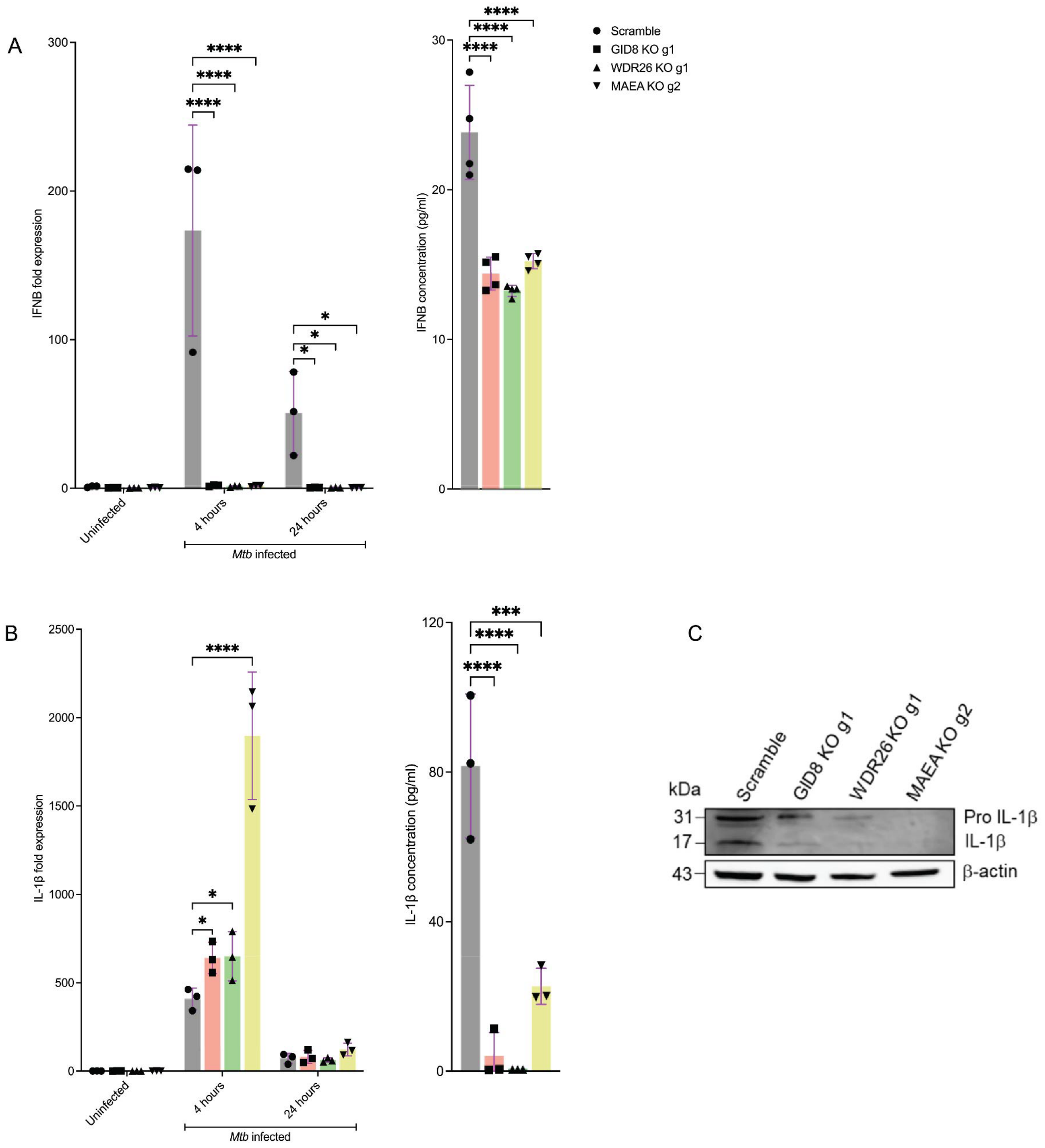


**Fig. S8. Related to Fig.5**

(A) Analysis of the expression of interferon 1 beta (IFNB) in scramble or *GID8*, *MAEA* and *WDR26* knockdown *Mtb* infected macrophages. hBMDMs were infected at MOI 4 and RNA was extracted for qPCR at 4 and 24 hours post infection. ELISA was carried out in culture supernatants 24 hours post infection; n=3. *P < 0.05; ****P < 0.0001.

(B) Analysis of the expression of IL-1β in scramble or *GID8*, *MAEA* and *WDR26* knockdown *Mtb* infected macrophages. qPCR and ELISA assays were carried out as in A. n=3 *, $P < 0.05$; ***P < 0.001; ****, $P < 0.0001$.

(D) Western blot analysis of IL-1β in scramble or *GID8*, *MAEA* and *WDR26* knockdown *Mtb* infected macrophages. Protein lysates were prepared from scramble or mutant hBMDMs infected with *Mtb* at MOI 4 for 24 hours
